# Translational initiation factor eIF5 replaces eIF1 on the 40S ribosomal subunit to promote start-codon recognition

**DOI:** 10.1101/366500

**Authors:** Jose L. Llácer, Tanweer Hussain, Adesh K. Saini, Jagpreet Nanda, Sukhvir Kaur, Yuliya Gordiyenko, Rakesh Kumar, Alan G. Hinnebusch, Jon R. Lorsch, V. Ramakrishnan

## Abstract

In eukaryotic translation initiation AUG recognition of the mRNA requires accommodation of Met-tRNA_i_ in a “P_IN_” state, which is antagonized by the factor eIF1. eIF5 is a GTPase activating protein (GAP) of eIF2 that additionally promotes stringent AUG selection, but the molecular basis of its dual function was unknown. We present a cryo-electron microscopy (cryo-EM) reconstruction of a 48S pre-initiation complex (PIC), at an overall resolution of 3.0 Å, featuring the N-terminal domain (NTD) of eIF5 bound to the 40S subunit at the location vacated by eIF1. eIF5 interacts with and allows a more accommodated orientation of Met-tRNA_i_. Substitutions of eIF5 residues involved in the eIF5-NTD/tRNA_i_ interaction influenced initiation at near-cognate UUG codons *in vivo*, and the closed/open PIC conformation *in vitro*, consistent with direct stabilization of the codon:anticodon duplex by the wild-type eIF5-NTD. The present structure reveals the basis for a key role of eIF5 in start-codon selection.

## Introduction

Eukaryotic translation initiation is a multistep process that involves assembly of a pre-initiation complex (PIC) comprised of the small (40S) ribosomal subunit, methionyl initiator tRNA (Met-tRNA_i_) and numerous eukaryotic initiation factors (eIFs). The binding of this 43S PIC to the capped 5ʹ end of mRNA is followed by scanning the mRNA leader for the correct AUG start codon. The binding of eIF1 and eIF1A to the 40S subunit promotes a scanning-conducive, open conformation favourable for rapid binding of Met-tRNA_i_ as a ternary complex (TC) with eIF2-GTP, in a conformation (P_OUT_) suitable for scanning successive triplets in the 40S P site for complementarity to the anticodon of Met-tRNA_i_. The multisubunit eIF3 complex also binds directly to the 40S subunit and stimulates 43S assembly, attachment to mRNA, and subsequent scanning. During the scanning process, hydrolysis of GTP in TC is stimulated by the GTPase activating protein (GAP) eIF5, but release of phosphate (P_i_) from eIF2-GDP-P_i_ is prevented by the gatekeeper molecule eIF1 at non-AUG codons. Recognition of an AUG start codon induces a major conformational change in the PIC to a scanning-arrested closed (P_IN_) complex made possible by the dissociation of eIF1, which eliminates a clash it would have with Met-tRNA_i_ in its fully accommodated P_IN_ conformation. The change is accompanied by the movement of the C-terminal tail (CTT) of eIF1A from eIF1 towards the GAP domain of eIF5, an event that triggers P_i_ release from eIF2. The P_IN_ conformation is further stabilized by direct interaction of the unstructured N-terminal tail (NTT) of eIF1A with the codon-anticodon duplex (Hinnebusch, 2014; Hinnebusch, 2017; Aylett and Ban, 2017).

Biochemical and genetic analysis of the effects of mutations in different eIFs on stringent selection of AUG start codons *in vivo* and on partial initiation reactions in a reconstituted yeast translation system have helped to reveal their molecular functions in controlling the transition between the open/P_OUT_ and closed/P_IN_ states of the scanning PIC. For example, mutations in eIF1 that weaken its binding to the 40S subunit allow the transition from open/P_OUT_ to closed/P_IN_ states to occur more frequently at near-cognate UUG codons, conferring the Sui^−^ (*<u>su</u>*ppressor of *<u>i</u>*nitiation codon) phenotype *in vivo* (Cheung et al., 2007) (Martin-Marcos et al., 2013). By contrast, eIF1 substitutions that increase its affinity for the ribosome and decrease its rate of dissociation from the PIC — thus impeding the open/P_OUT_ to closed/P_IN_ transition — confer the Ssu^−^ (*s*uppressor of Sui^−^) phenotype by suppressing elevated UUG initiation conferred by other Sui^−^ mutations (Martin-Marcos et al., 2014). Ssu^−^ substitutions in the eIF1A NTT were also shown to decrease the rate of eIF1 dissociation, and to destabilize the closed conformation of the PIC (Fekete et al., 2007; Cheung et al., 2007).

In addition to its function as a GAP for eIF2, eIF5 has been implicated in stringent selection of AUG start codons. eIF5 consists of an N-terminal domain (NTD, 1-170 residues), which is connected by a long flexible linker to a C-terminal domain (CTD) (Conte et al., 2006) (Wei et al., 2006). The eIF5-NTD contains the GAP function of eIF5, which is stimulated by the PIC and requires the conserved Arg-15 located in its unstructured N-terminus, possibly functioning as an “arginine finger” that interacts with the GTP binding pocket in eIF2γ to stabilize the transition state for GTP hydrolysis (Algire et al., 2005) (Das et al., 2001) (Paulin et al., 2001). Substitution of Gly-31 with arginine in the eIF5-NTD (the *SUI5* allele) is lethal but confers a dominant Sui^−^ phenotype in yeast cells (Huang et al., 1997). Consistent with this, in the reconstituted system, the eIF5-G31R substitution alters the regulation of P_i_ release such that it occurs faster at UUG versus AUG start codons; and it modifies a functional interaction of eIF5 with the eIF1A-CTT to favour the closed PIC conformation at UUG over AUG start codons. Biochemical analysis of intragenic Ssu^−^ suppressors of *SUI5* indicated that the effect of eIF5-G31R in both deregulation of P_i_ release and partitioning of PICs between open and closed states contribute to the enhanced UUG initiation *in vivo* (Maag et al., 2006) (Saini et al., 2014). Because the location of eIF5 in the PIC was unknown, it has been unclear how the G31R substitution alters these events in the open/P_OUT_ to closed/P_IN_ transition at the molecular level. Movement of the wild-type eIF5-NTD and the eIF1A CTT towards one another within the PIC is triggered by AUG recognition, and this rearrangement is dependent on scanning enhancer (SE) elements in the eIF1A-CTT (Nanda et al., 2013). The fact that mutations in SE elements more strongly reduced the rate of P_i_ release than eIF1 dissociation suggested that the SE-dependent movement of the eIF5-NTD towards the eIF1A-CTT facilitates P_i_ release following eIF1 dissociation (Nanda et al., 2013).

The eIF5 C-terminal domain (CTD) also performs multiple functions in assembly of the PIC and control of start codon selection such as promoting the closed PIC conformation by enhancing eIF1 dissociation (Nanda et al., 2009; Nanda et al., 2013). Accordingly, overexpressing eIF5 in yeast (Nanda et al., 2009) or mammalian cells (Loughran et al., 2012) relaxes the stringency of start codon selection presumably by enhancing eIF1 release (Loughran et al., 2012). As might be expected, overexpressing eIF1 has the opposite effect of increasing discrimination against near-cognates in both yeast (Valasek et al., 2004) (Fekete et al., 2007) and mammals (Ivanov et al., 2010). The opposing effects of eIF1 and eIF5 on start codon stringency are exploited to negatively autoregulate their own translation, and through cross-regulation of each other, maintain balanced expression of the two proteins and stabilize the stringency of AUG selection (Loughran et al., 2012). eIF5 function is further regulated by an inhibitory protein (Tang et al., 2017) (Kozel et al., 2016). Thus, the structure of a eukaryotic translation initiation complex containing eIF5 would greatly aid our understanding of its multiple functions.

The recent structures of eukaryotic translation initiation complexes from yeast (Aylett et al., 2015) (Hussain et al., 2014) (Llácer et al., 2015) as well as mammals (Lomakin and Steitz, 2013) (des Georges et al., 2015) (Hashem et al., 2013) (Simonetti et al., 2016) have provided many insights into the molecular events involved in scanning and AUG recognition. eIF1, eIF1A, TC and eIF3 have been observed in an open (P_OUT_) conformation, as well as a scanning-arrested closed (P_IN_) conformation of the 40S (Simonetti et al., 2016) (Llácer et al., 2015). In a partial 48S pre-initiation complex from yeast (py48S), we had tentatively suggested that an unassigned density at low resolution near eIF2γ belongs to eIF5-CTD (Hussain et al., 2014). However, there is no clear structural information on the position and conformation of either the CTD or NTD of eIF5 in the PIC.

Here we have determined a cryo-EM structure of a yeast 48S complex in the P_IN_ conformation at near atomic resolutions (3.0 Å to 3.5 Å maps) containing clear densdity for the eIF5-NTD (py48S-eIF5N). Remarkably, in this py48S-eIF5N complex, eIF1 has been replaced by the eIF5-NTD, which is bound near the P-site at essentially the same position eIF1 binds in the open/P_IN_ state. The tRNA_i_ is more fully accommodated in the P site than observed in previous structures containing eIF1, and is also tilted toward the 40S body, apparently setting the stage for its interaction with eIF5B and subsequent joining of the 60S subunit. Extensive interaction with the eIF5-NTD appears to stabilize this tRNA_i_ conformation, using two β-hairpins structurally analogous to those in eIF1 that oppose tRNA_i_ accommodation in the scanning complex. Mutations expected to weaken the observed eIF5-NTD/tRNA_i_ interactions diminish initiation at near-cognate UUG codons *in vivo* and disfavour transition to the closed/P_IN_ conformation at UUG codons in reconstituted PICs *in vitro*, whereas mutations expected to stabilize the interactions have the opposite effects. These and other results suggest that the eIF5-NTD stabilizes the codon-anticodon interaction and the closed/P_IN_ state of the PIC, prevents eIF1 rebinding, and promotes a conformation of the 48S PIC compatible with eIF5B binding and subunit joining.

## RESULTS

### Overview of the cryo-EM structure of a yeast 48S PIC containing eIF5

A partial yeast 48S PIC containing clear density for eIF5-NTD (py48S-eIF5N) was reconstituted by sequential addition of purified *Saccharomyces cerevisiae* eIF1, eIF1A, eIF3, eIF5, TC and an eIF4F-eIF4B-mRNA complex to 40S subunits purified from the yeast *Kluyveromyces lactis*. An unstructured, capped 49-mer mRNA with an AUG codon and an optimal Kozak sequence for yeast was used (See Experimental Procedures). Native eIF2, eIF3 and tRNA_i_ were purified from *Saccharomyces cerevisiae*, whereas all other eIFs were produced in bacteria. In our earlier studies, eIF4 factors were not used to deliver the mRNA to the PIC, and the mRNA was uncapped and lacked an optimal Kozak sequence. These changes in assembly protocol may have helped in capturing eIF5 in the 48S.

The structure of py48S-eIF5N was determined to an overall resolution of 3.0 Å to 3.5 Å in respective maps: 1, A, B, C1 and C2 (Figure 1 – figure supplement 1, 2 and 3; Tables 1, 2 and 3), and the resulting model is shown in Figure 1. The local resolution and density are greatest for the 40S core and ligands directly attached to it, including the eIF5-NTD (Figure 1 – figure supplement 4 and Table 2). Met-tRNA_i_ is bound to py48S-eIF5N in a P_IN_ state (base paired to the AUG codon), with a closed mRNA latch and compressed conformation of h28 (the rRNA helix connecting the 40S head and body) (Figure 1 – figure supplement 5), as in previous py48S complexes exhibiting a closed conformation of the 40S, and in contrast to the open-latch and relaxed h28 conformations of a py48S open complex (Figure 1 – figure supplement 5) (Hussain et al., 2014) (Llácer et al., 2015). Density for the eIF5-NTD is observed on the 40S platform near the P-site (Figure 2A), which is also the site for binding of eIF1 (Llácer et al., 2015) (Hussain et al., 2014) (Lomakin and Steitz, 2013) (Hashem et al., 2013) (Rabl et al., 2011) (Aylett et al., 2015). Although eIF1 and the eIF5-NTD share structural similarity (Conte et al., 2006) (Figure 2 – figure supplement 1A-B), the resolution of the map allowed us to unambiguously determine that the density belongs to eIF5-NTD and not eIF1 (See below for details of fitting eIF5-NTD into the density map). Hence, py48S-eIF5N represents a later stage in eukaryotic translation initiation after the recognition of the start codon and dissociation of eIF1 but prior to dissociation of eIF2 from Met-tRNA_i_. The latter was expected because nonhydrolyzable GDPCP was used instead of GTP in assembly of TC; and GTP hydrolysis is required to reduce the affinity of eIF2 for Met-tRNA_i_ (Kapp and Lorsch, 2004). Clear density for the codon:anticodon interaction is observed at the P-site; and the complete N-terminal tail (NTT) of eIF1A was resolved, stabilizing the codon:anticodon helix as seen in earlier closed py48S complexes containing eIF1 instead of eIF5-NTD (Llácer et al., 2015) (Hussain et al., 2014).

**Figure 1.**
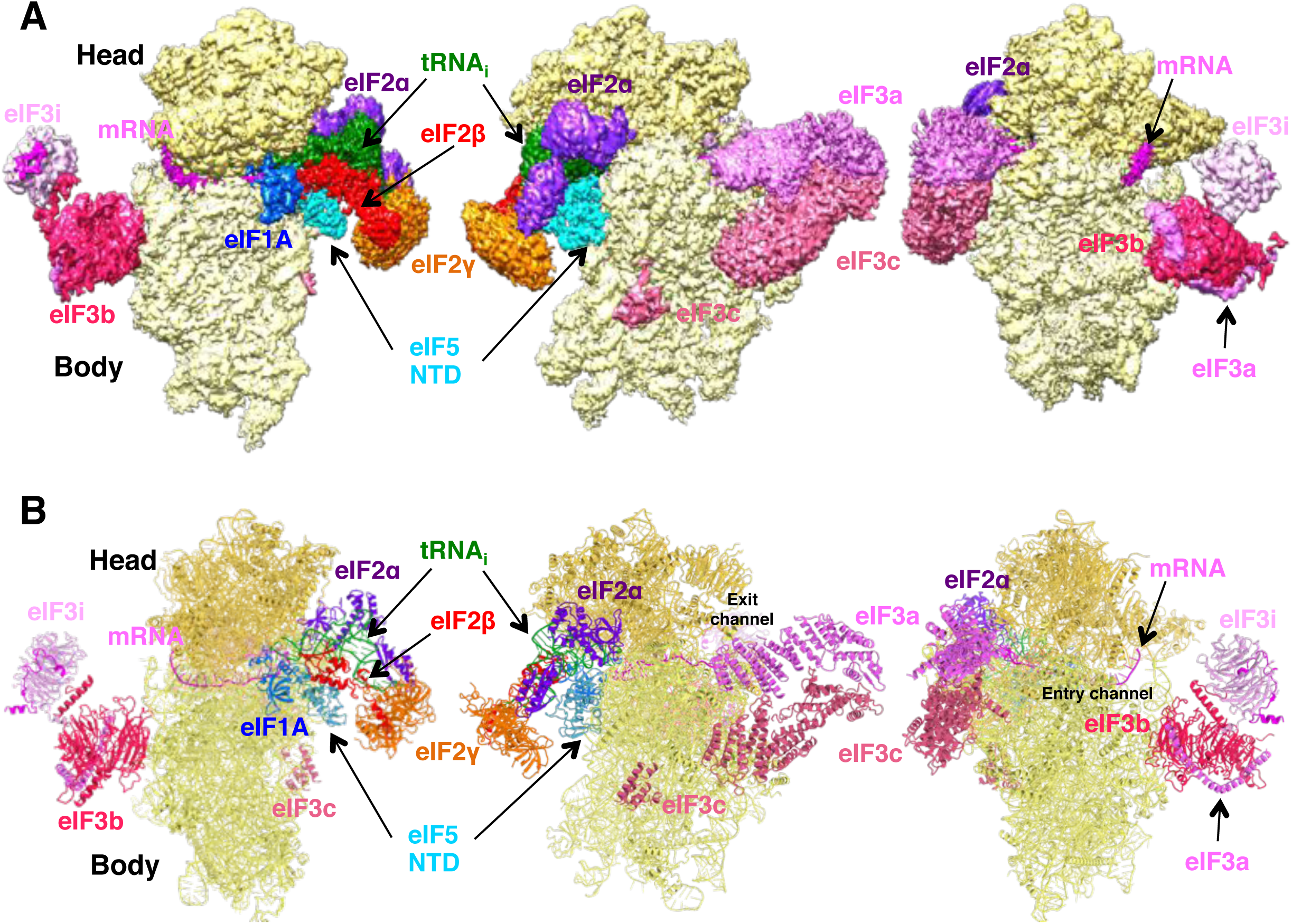
Cryo-EM structure of py48S-eIF5N. (A) CryoEM maps of the PIC py48S-eIF5N shown in three orientations. Regions of the map are colored by component to show the 40S subunit (yellow), eIF1A (blue), eIF5-NTD (cyan), Met-tRNA_i_^Met^ (green), mRNA (magenta), eIF2α (violet), eIF2γ (orange), eIF2β (red), eIF3 (different shades of pink). The 40S head is shown in a darker yellow compared to the body. The density for 40S, eIF1A, mRNA, tRNA, eIF2 subunits and eIF5 is taken from Map C1, whereas density for eIF3 PCI domains is taken from Map A, and for eIF3-bgi subcomplex from Map B. The same colors are used in all the figures. (B) Atomic model for the PIC in the same colors and in the same three orientations. See also Figure 1 – figure supplement 1, 2, 3, 4, 5.

In Map 1, the occupancies for eIF3, eIF2γ, eIF2β and the acceptor arm of the tRNA_i_ are rather low. In order to observe the location of these ligands on the py48S-eIF5N, a larger data set and extensive 3D classification using different masks was employed to obtain multiple py48S-eIF5N maps (Maps A, B, C1 and C2) showing clear densities for eIF1A, eIF3, TC (including eIF2β), eIF5 and mRNA (Figure 1 – figure supplement 1; See Supplemental Experimental Procedures). Density for the mRNA is present throughout the mRNA channel from the entry to the exit site; the mRNA interacts with the 40S head at the entry site and with the a-subunit of eIF3 (eIF3a) and other ribosomal elements at the exit site (Figure 1B). The density for different subassemblies of eIF3 subunits (in maps A and B), including the dimer of eIF3a/eIF3c PIC domains and the β-propellers of eIF3b and eIF3i, are observed at the solvent side of the 40S. Only density corresponding to the helical bundle formed by the eIF3c-NTD is observed at the subunit interface (Figure 1A-B). Densities corresponding to the eIF5-CTD and eIF4 factors were not observed in any of these py48S-eIF5N maps, which presumably reflects the flexibility or dynamic nature of these domains/factors in the 48S PIC during the later steps of initiation.

### The eIF5-NTD replaces eIF1 on the 40S platform near the P-site

A clear and distinct density for an ‘eIF1-like’ domain was observed at the top of h44 near the P-site (Figure 2A), but fitting the known eIF1 structure into this density as seen in previous py48S-maps (Hussain et al., 2014) (Llácer et al., 2015) could not account for all of it (Figure 2B). Also, there was no density to account for the C-terminal β-strand of eIF1. Moreover, close inspection revealed discrepancies between the densities and side chains of eIF1, particularly for β-hairpin 1 at the P-site. Together, the known structural similarity of the eIF5-NTD with eIF1 (PDB: 2E9H, (Conte et al., 2006)) (Figure 2 – figure supplement 1A-B), our previous suggestion that the eIF5-NTD might occupy the position of eIF1 on the 40S following eIF1 dissociation (Nanda et al., 2009), and our previous demonstration that the eIF5-NTD can bind directly to the 40S subunit (Nanda et al., 2013), all prompted us to place the NMR structure of eIF5-NTD from *Homo sapiens* into the unassigned density on the 40S platform. The eIF5-NTD structure accounted for the entire density, including the zinc binding domain (ZBD) absent in eIF1 (Figure 2B), and the high resolution of the map enabled us to unambiguously model the eIF5-NTD at the atomic level (Figure 1 – figure supplement 3C).

**Figure 2.**
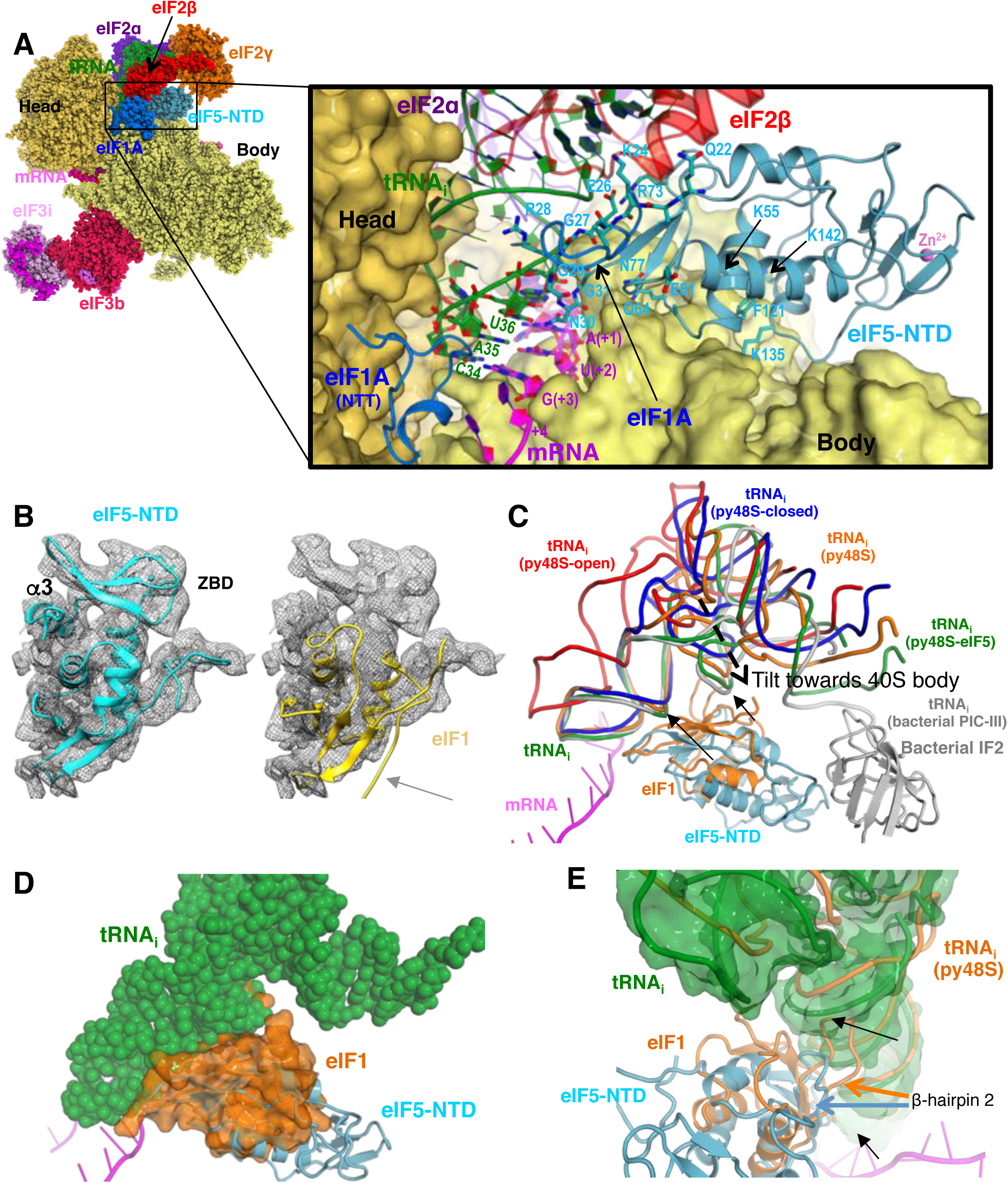
Contacts of eIF5-NTD with the other components in the 48S PIC. (A) A detailed view of the contacts of eIF5-NTD near the P site with the 40S subunit, tRNA_i_, mRNA, eIF1A and eIF2β. eIF5 residues involved in the contacts are shown in sticks. (B) Fitting of eIF5-NTD (left) and eIF1(right) on density in Map 1(low-pass filtered to 5Å). Zinc binding domain (ZBD) and helix α3 of eIF5, both absent in eIF1, are labeled. β5 of eIF1, which is not present in eIF5 is highlighted by a grey arrow. (C) The relative movement of the initiator tRNA in all reported yeast PICs, as deduced by a superposition using the 40S body. tRNA_i_s from py48S-eIF5N (this study; green), py48S PIC (PDB 3J81; orange), py48S PIC-closed (PDB 3JAP; blue) and py48S PIC-open (PDB 3JAQ red) are shown. eIF5-NTD from py48S-eIF5N and eIF1 from py48S PIC (PDB 3J81) are also shown. For comparison, tRNA_i_ and IF2 from a bacterial PIC with accommodated P site tRNA conformation is also shown (PICIII; PDB 5LMV; grey). In all closed conformations, the tip of the ASL is essentially in the same position; however, there is a different tilting of the tRNA_i_ towards the 40S body in the different PICs. eIF1 would clash with tRNA_i_ in py48S-eIF5N; black arrows highlight these clashes. (D) Representation of how eIF1 in py48S (transparent orange surface) would clash with tRNA_i_ in py48S-eIF5N (in spheres). The model results from aligning the 40S bodies of the two structures. See also Figure 2 – figure supplement 1, 2. (E) eIF1 and eIF5-NTD share a similar fold, however β-hairpin 2 in eIF5 is shorter than that in eIF1, which allows a further accommodation of tRNA_i_ in the P site. eIF1 and tRNA_i_ from py48S (PDB 3J81; orange) are superimposed on eIF5-NTD/tRNA_i_ from py48S-eIF5N.

The eIF5-NTD binds on the platform at essentially the same location occupied by eIF1 in previous py48S structures (Figures 2A,C and Figure 2 – figure supplement 1C-F and Movie 1), interacting with 18S rRNA residues in h44 (1760; *S. cerevisiae* numbering), h45 (1780 and 1781) and h24 (994, 995, 1001, 1002 and 1004). In this position, eIF5-NTD interacts with eIF1A, as does eIF1 in other py48S structures; and also makes limited contacts with eIF2β and eIF2γ (Figure 2 – figure supplement 1C). However, residue Arg15 of eIF5-NTD (essential for its GAP activity; (Algire et al., 2005)) is positioned more than 10 Å away from the bound GTP analog in eIF2γ (Figure 2 – figure supplement 1D). Accordingly, this position and conformation of the eIF5-NTD does not appear compatible with the GAP activity of eIF5 GAP and likely instead represents a state after release of inorganic phosphate (P_i_) would normally have occurred. Multiple residues in the eIF5-NTD, including Lys24, Gly27, Arg28, Gly29, Asn30, and Gly31 (in β-hairpin 1), and Lys71 and Arg73 (in β-hairpin 2), make multiple contacts with the anticodon stem loop (ASL) of the tRNA_i_ (Figure 2A, Figure 2 – figure supplement 1C) and these contacts are more extensive and more favorable than are those made by the structurally analogous β-hairpins 1 and 2 of eIF1 in py48S (Figure Figure 2 – figure supplement 1E,F and Movie 1) (Hussain et al., 2014). Interestingly, β-hairpin 1 of eIF5-NTD is positioned in the mRNA channel at the P-site and monitors the codon:anticodon interaction in a similar fashion as the β-hairpin 1 of eIF1 (Figure 2A,C)(Hussain et al., 2014) (Martin-Marcos et al., 2013). The conserved Asn30 in β-hairpin 1 of eIF5 makes contacts with both the codon and anticodon (Figure 2A,C). However, in eIF5-NTD the β-hairpin 1 is shorter and contains three Gly residues compared to only one Gly in the case of eIF1. Gly27, Gly29 and Gly31 are closely packed against the ASL (Figure 2A) and any larger residue would create a steric clash with the ASL. The β-hairpin 2 of eIF5-NTD is also three residues shorter than that of eIF1 and is oriented away from the tRNA_i_ to allow the latter to be tilted more towards the 40S body compared to previous py48S complexes that contain eIF1 (Hussain et al., 2014) (Figure 2C-E and Movie 1).

A superimposition of this structure with the previous py48S structure containing eIF1 instead of the eIF5-NTD (Hussain et al., 2014) reveals that eIF1 would sterically clash with tRNA_i_ at its position in py48S-eIF5N (Figures 2D-E and Movie 1) indicating a further accommodation of the tRNA_i_ in the P site after dissociation of eIF1. In fact, the tRNA_i_ conformation in the current structure most closely resembles that found in bacterial translation initiation complexes (Hussain et al., 2016). Of the various eukaryotic py48S complexes containing eIF2, the structure here shows the maximum degree of tRNA_i_ accommodation and tilt towards the 40S body (Figure 2C). This tRNA_i_ tilt towards the body is also similar to that found in eukaryotic 80S initiation complexes containing eIF5B but lacking eIF2, eIF1 and eIF5 (Figure 2 – figure supplement 2A) (Fernandez et al., 2013; Yamamoto et al., 2014). Given that the affinity of TC for the PIC increases when eIF1 is ejected after AUG recognition (Passmore et al., 2007) (Nanda et al., 2013), the tRNA_i_ conformation observed in the present complex probably represents its most stable conformation, and it is conceivable that eIF5-NTD participates in this stabilization via its interaction with the ASL.

### eIF5-NTD substitutions at the codon:anticodon interface alter the stringency of AUG start codon selection *in vivo*

To examine the physiological significance of the direct contacts observed between the eIF5-NTD and the start codon and ASL, specific residues were selected for mutagenesis based on their proximity to tRNA_i_ or 18S rRNA in the PIC (Figure 3A-B). Mutations were generated in the gene coding for eIF5 harboring a C-terminal FLAG epitope (the *TIF5-FL* allele), on a *LEU2* plasmid, and introduced into a *his4-301 tif5Δ* strain lacking chromosomal *TIF5* and carrying WT *TIF5* on a *URA3* vector. The *his4-301* mutation confers auxotrophy for histidine (His^−^) owing to the absence of the AUG start codon of the WT *HIS4* allele, which can be suppressed by Sui^−^ mutations that allow utilization of the third, UUG triplet as start codon, including the eIF1 mutation *sui1-L96P* (Martin-Marcos et al., 2011) (Figure Figure 3 – figure supplement 1A). Hence, by determining effects of the eIF5 mutations on the histidine requirement of the resulting *his4-301* strains, we could determine their effects on accurate start codon recognition *in vivo*.

**Figure 3.**
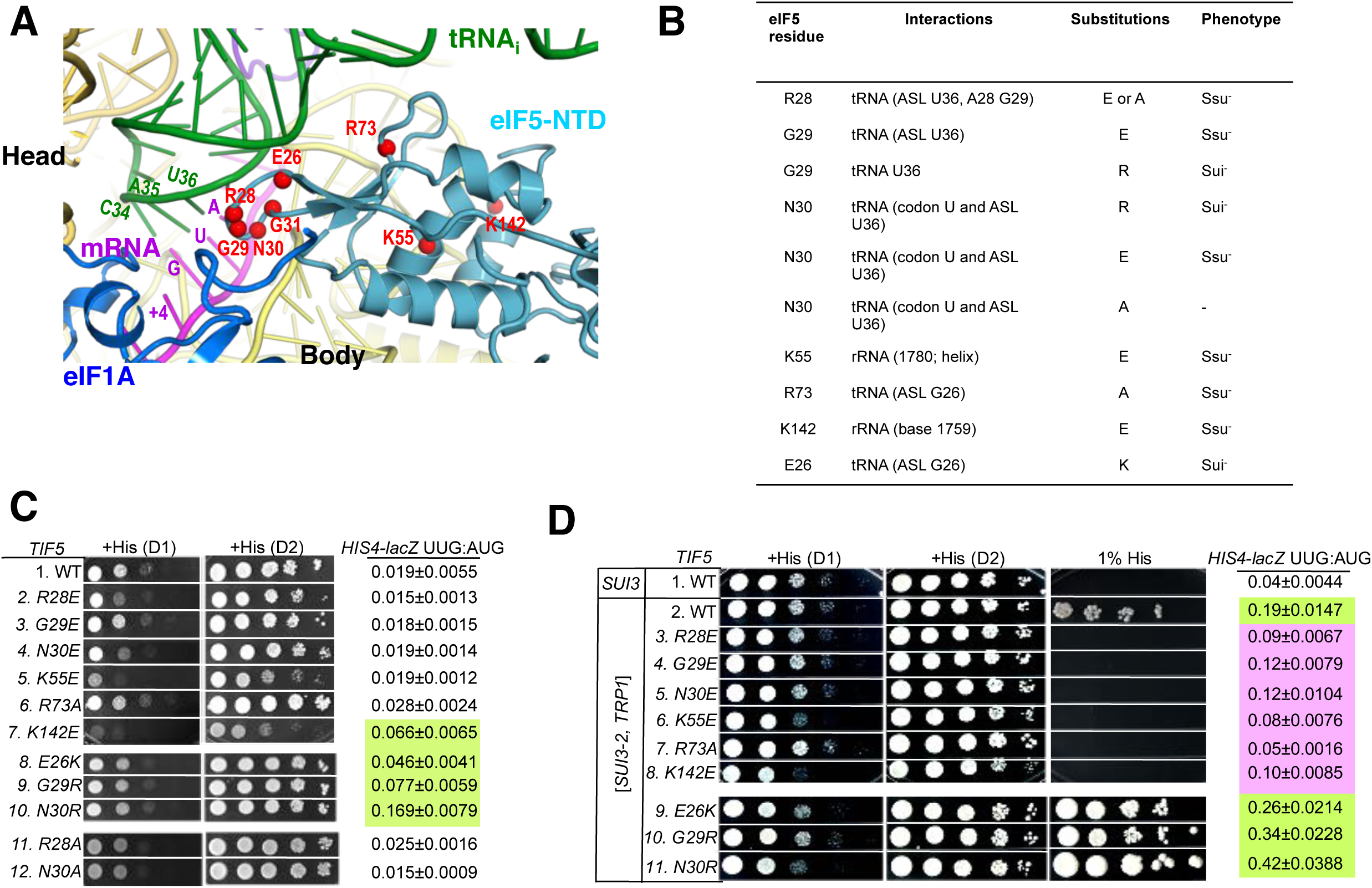
Genetic evidence that contacts of the eIF5-NTD with the tRNA_i_ ASL and mRNA AUG codon in py48S-eIF5N are crucial for stringency of start codon recognition in vivo. (A) Location of the eIF5 residues substituted in genetic studies, highlighted in red and shown as spheres. (B) Summary of eIF5-NTD residues substituted by *TIF5* mutations (col. 1), their interactions in py48S-eIF5N (col. 2), the amino acid substitutions introduced (col. 3), and the observed Sui^−^ or ++Ssu^−^ phenotypes in vivo (col. 4) revealed by results in (C or D). (C) Slg^−^ and His^+^/Sui^−^ phenotypes were determined for derivatives of *his4-301 tif5Δ* strain ASY100 harboring the indicated *TIF5-FL* alleles on *LEU2* plasmids. Ten-fold serial dilutions of the strains were spotted on synthetic complete medium lacking leucine (SC-L) and supplemented with 0.3 mM histidine (+His), and incubated for 1d (D1) or 2d (D2) at 30°C. To assess the effect of *TIF5* substitutions on start codon recognition, the strains were transformed with *HIS4-lacZ* reporter plasmids containing AUG (p367) or UUG (p391) start codons. Cells were cultured in SC medium lacking leucine and uracil at 30°C and β-galactosidase activities were measured in whole cell extracts (WCEs). Ratios of β-galactosidase expressed from the UUG to AUG reporter were calculated from four independent transformants and mean ratios and S.E.M.s (error bars) are reported on the right. Ratios indicating Sui^−^ phenotypes are highlighted in lime green. (D) Strains described in (C) were transformed with *SUI3-2* plasmid pRSSUI3-S264Y-W (rows 2–11) or empty vector (row 1). Slg^−^ and His^+^/Sui^−^ phenotypes were determined by spotting the 10-fold serial dilutions of strains on SC medium lacking leucine and tryptophan and supplemented with either 0.3 mM histidine (+His) or 0.0003 mM histidine (1% His), and incubated for 1d (D1) or 2d (D2) for +His medium and 6d for 1% His medium, at 30°C. The *HIS4-lacZ* initiation ratios, were determined as in (B) except the cells were grown in SC medium lacking leucine, uracil and tryptophan. Ratios indicating Sui^−^ (lime green) or Ssu^−^ (pink) phenotypes are highlighted. See also Figure 3 – figure supplement 1

After plasmid-shuffling to evict WT *TIF5*, we found that all mutant strains were viable, but that the strains carrying *TIF5-FL* alleles -*R28E, -N30E, -K55E*, and -*K142E* displayed slow-growth (Slg^−^) phenotypes of varying degrees compared to the WT *TIF5-FL* strain (Figure 3C; +His (D1)). None of the mutations conferred any marked differences in steady-state expression of the FLAG-tagged eIF5 proteins (Figure 3 – figure supplement 1B,C). None of the mutants exhibited a His^+^ phenotype on media lacking His or containing only 1% of the His used to fully supplement the auxotrophy (data not shown), suggesting the absence of marked Sui^−^ phenotypes. However, assaying β-galactosidase expressed from matched *HIS4-lacZ* reporters with either AUG or UUG start codons revealed that *E26K*, *K142E*, *G29R* and *N30R* conferred ~2-, 3-, 4-and 8-fold increases in the UUG:AUG ratio, respectively (Figure 3C, *HIS4-lacZ* UUG:AUG, rows 7-10 vs. 1). The absence of a His^+^ phenotype for these mutations might result from a failure to increase the UUG:AUG ratio above a critical threshold level (Martin-Marcos et al., 2013); the strong Slg^−^ phenotype of *K142E* would likely also impede growth on –His or 1% His medium.

Interestingly, the structure reveals that residues E26, G29 and N30 are in proximity to the anticodon stem loop (ASL) of tRNA_i_ (Figure 3A-B), and we hypothesized that increasing the positive charge of these residues by replacing them with R or K might stabilize the P_IN_ state of TC binding even on near-cognate UUG start codons, thus accounting for the increased UUG:AUG initiation ratios conferred by the E26K, G29R, and N30R substitutions (Figure 3C). If so, then decreasing the positive charge of residues 28 and 73, which also approach the tRNA_i_ ASL, by the *R28A, R28E*, and *R73A* mutations, might destabilize the P_IN_ state and increase discrimination against UUG start codons. To test this idea, the *TIF5-FL* strains were transformed with plasmid-borne *SUI3-2*, encoding the S264Y substitution in eIF2β that confers a dominant His^+^/Sui^−^ phenotype and elevates the UUG:AUG ratio in otherwise WT strains (Huang et al., 1997).

As expected, *SUI3-2* confers growth on media containing 1% His and elevates the UUG:AUG ratio by ~5-fold (Figure 3D, *HIS4-lacZ* UUG:AUG rows 1-2). Importantly, the His^+^/Sui^−^ phenotype of *SUI3-2* is suppressed efficiently by the *-R28E, -R28A*, and *-R73A* alleles of *TIF5-FL*, which also substantially diminish the UUG:AUG ratio in *SUI3-2* cells (Figure 3D, rows 3 & 7 vs. 2; Figure 3 – figure supplement 1E, rows 2 & 4), thus conferring <u>S</u>uppressor of <u>Sui</u>^−^ (Ssu^−^) hyperaccuracy phenotypes. Interestingly, *G29E, N30E, K55E* and *K142E* also suppress the His^+^/Sui^−^ phenotype and mitigate the elevated UUG:AUG ratio conferred by *SUI3-2* (Figure 3D, rows 4-6, 8 vs. 2). The Ssu^−^ phenotypes of *R28E*, *N30E* and *K55E* are dominant in strains harboring *SUI3-2* and WT *TIF5* (Figure 3 – figure supplement 1D), suggesting that these eIF5 variants can efficiently compete with WT eIF5 for incorporation into PICs but function poorly in stabilizing the P_IN_ state at UUG codons.

The *TIF5-FL* alleles *E26K, G29R* and *N30R*, which elevate the UUG:AUG ratio in otherwise WT cells (Figure 3C, rows 7-10 vs. 1), also exacerbate the His^+^/Sui^−^ defect of *SUI3-2* (Figure 3D, 1% His, rows 9-11 vs. 2) and confer ~1.4-, 1.8-and 2.2-fold increases in the UUG:AUG ratio compared to *SUI3-2* cells containing WT *TIF5-FL* (Figure 3D, *HIS4*-*lacZ* UUG:AUG rows 9-11 vs. 2). *TIF5-FL-N30R* also conferred a modest Slg^−^ phenotype in *SUI3-2* cells (Figure 3D, +His, row 11 vs. 2), which was not seen in otherwise WT cells containing this allele (Figure 3C, row 10 vs. 1). The ability of *N30R* and *E26K* to intensify the His^+^/Sui^−^ defect of *SUI3-2* is dominant, occurring in strains harboring WT *TIF5* (Figure 3 – figure supplement 1C, rows 6-7 vs. 2), indicating that these eIF5 variants can also compete with WT eIF5 for incorporation into the PIC and stabilize the P_IN_ state at UUG codons.

It is noteworthy that substitutions of G29 introducing positively or negatively charged residues evoke opposing phenotypes (Figure 3C-D), with G29R conferring increased UUG initiation in otherwise WT cells and exacerbating the Sui^−^ phenotype of *SUI3-2*; and G29E suppressing the Sui^−^ defects of *SUI3-2*, an Ssu^−^ phenotype (Figure 3C-D, *G29R, G29E*). These findings are consistent with the possibility that introducing a positive charge at this position of eIF5 at the interface with tRNA_i_ enhances electrostatic attraction with the ASL to stabilize the P_IN_ state at UUG start codons; whereas a negatively charged side-chain at this position destabilizes P_IN_ through electrostatic repulsion with the ASL to preferentially diminish selection of UUG codons, which form mismatched duplexes with the tRNA_i_ ASL. The same reasoning can explain the opposing phenotypes of the Arg (Sui^−^) and Glu (Ssu^−^) substitutions of N30 (Figure 3C-D, *N30E*, *N30R*); and as noted above, the Ssu^−^ phenotype of R28E and Sui^−^ phenotype of E26K. The fact that R73A and R28A also confer Ssu^−^ phenotypes without introducing electrostatic repulsion underscores the importance of the native contacts of R28 and R73 with the tRNA_i_ ASL in stabilizing P_IN_. Presumably, the Ssu^−^ phenotypes of the K55E and K142E substitutions result from introducing electrostatic repulsion between these residues and 18S rRNA residues 1780 and 994/995, respectively, perturbing the eIF5-NTD:40S interface in a way that diminishes the formation or stability of the P_IN_ state at UUG codons. Together, the genetic data provide strong evidence that the contacts between the eIF5-NTD and the tRNA_i_ ASL visualized in the cryo-EM structure are crucial for a WT stringency of start codon recognition *in vivo*.

### eIF5-NTD substitutions at the codon:anticodon interface alter the influence of the start codon on transition to the closed PIC conformation

We have previously shown that monitoring dissociation of fluorescently labeled eIF1A from 48S PICs using fluorescence anisotropy is a useful tool to distinguish between *‘open’* (P_OUT_) and *‘closed’* (P_IN_) conformations of the PIC (Saini et al., 2014) (Maag et al., 2006). Although dissociation of eIF1A from the PIC at this stage of the initiation process is slow and does not appear to be a physiologically relevant event, it does report on the relative abundance and stability of the open and closed states of the complex. In WT PICs, dissociation of eIF1A occurs with biphasic kinetics, with the fast phase reflecting complexes in the open state, in which eIF1A is less stably bound, and the slow phase reflecting the more stable, closed state. The ratio of amplitudes of the slower phase (a_2_) over the fast phase (a_1_) is taken as the apparent equilibrium constant between the closed and open states (a_2_/a_1_) and is referred to as ‘K_amp_’. As observed in previous studies, with WT complexes K_amp_ is higher when the model mRNA has an AUG start codon (mRNA(AUG)) than when it has a near-cognate UUG codon (mRNA(UUG)) (5.9 vs. 3.2; Table 4A, rows 1-2), consistent with the closed state being more favored in the former case than in the latter. This effect is also reflected in the rate constants for the fast (k_1_) and slow (k_2_) phases, which are both higher for complexes assembled on mRNA(UUG) than on mRNA(AUG) (6 vs. 22×10^−3^ s^−1^ and 0.4 vs. 2.1×10^−3^ s^−1^, respectively; Table 4A and Figure 4A), indicating that eIF1A is less stably bound in both the open and closed states in complexes assembled on a near-cognate start codon. Consistent with this interpretation, the fluorescence anisotropy of the C-terminal fluorescein moiety on eIF1A is higher in complexes assembled on mRNA(AUG) (R_bound_ = 0.21) than on mRNA(UUG) (R_bound_ = 0.18) (Table 4A, rows 1-2). Because higher fluorescence anisotropies indicate less freedom of rotation of the fluorophore, these data indicate that a “tighter,” more constrained complex is preferentially formed on mRNA containing the cognate AUG codon.

**Figure 4.**
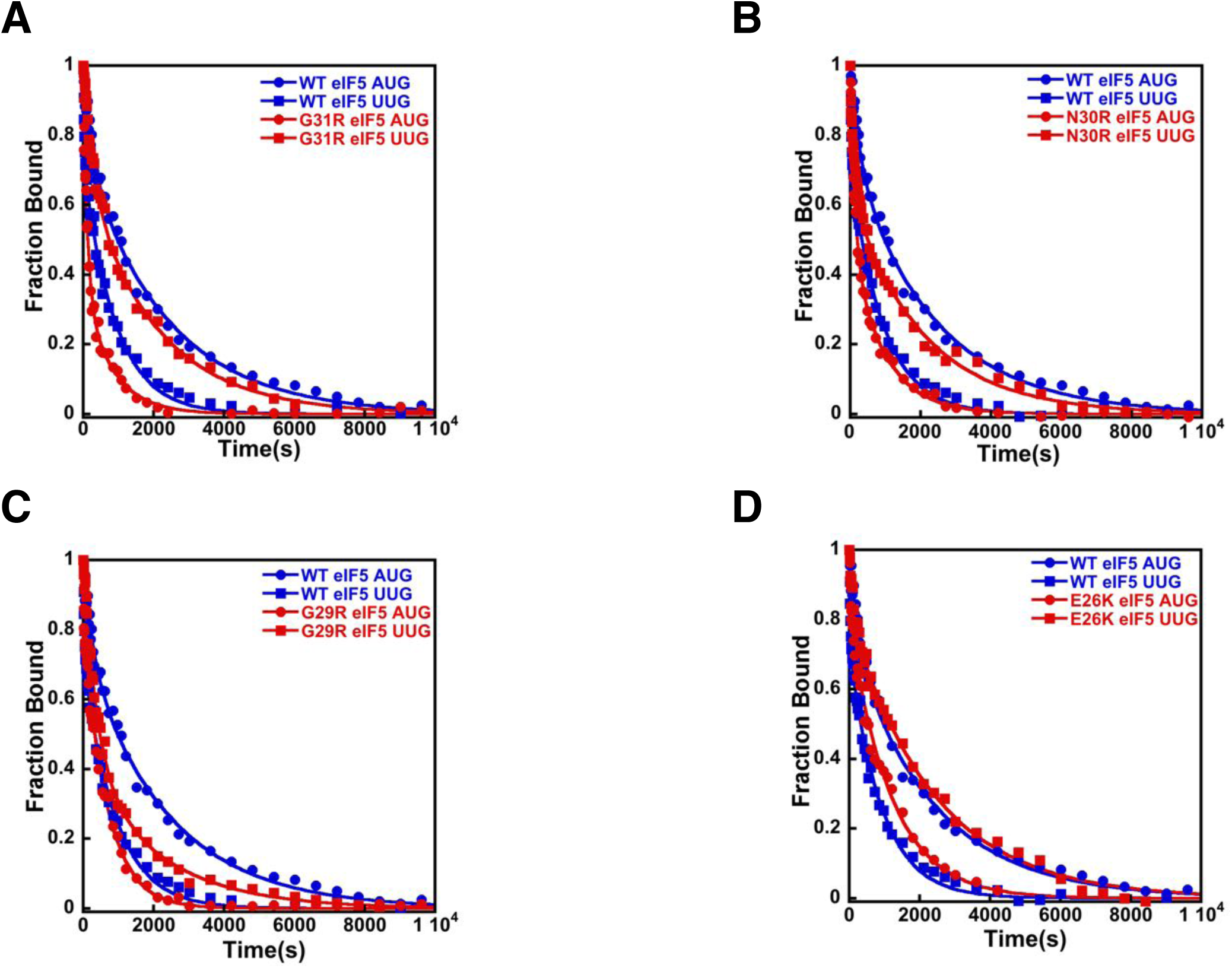
eIF5-NTD Sui^−^ mutants stabilize the closed conformation of the PIC at UUG codons. Dissociation of fluorescein tagged eIF1A from 43S·mRNA complexes reconstituted with model mRNAs containing an AUG or UUG start codon and either WT or Sui^−^ variants of eIF5 was monitored as decrease in fluorescence anisotropy over time after addition of excess unlabeled eIF1A. The data were fit with a double exponential decay equation. (A) eIF1A dissociation from 48S PICs assembled with WT eIF5 (blue) or eIF5-G31R (red) and mRNAs with an AUG (closed circles) or UUG (closed squares) start codons. (B-D) eIF1A dissociation from PICs containing WT eIF5 (blue) or the indicated mutant eIF5 (red) and mRNAs with AUG (closed circles) or UUG (closed squares) start codons, for eIF5-N30R (B), eIF5-G29R (C), or eIF5-E26K (D). Experiments were done at least 2 times and errors are calculated as average deviation

Using this system, we sought to determine the mechanistic impact of substitutions in the eIF5 residues that are in proximity to the start codon:tRNA_i_ anticodon helix. As described above, in the presence of WT eIF5, eIF1A is more stably bound to the PIC with mRNA(AUG) than mRNA(UUG) (Figure 4A blue closed circles versus blue closed squares) with a relatively higher K_amp_ for mRNA(AUG) (Table 4A; rows 1-2), indicating a greater preponderance of complexes in the closed state. As observed previously (Saini et al., 2014) (Maag et al., 2006), the G31R-eIF5 substitution, which has a strong, dominant Sui^−^ phenotype *in vivo*, inverts the effect of an AUG versus UUG start codon on the dissociation kinetics (Figure 4A red closed circles *versus* red closed squares). In contrast to WT eIF5, PICs containing G31R eIF5 have higher K_amp_ values at UUG than AUG start codons (Table 4A, rows 3-4), thus indicating the closed state of the PIC is favored in the former case relative to the latter, consistent with the Sui^−^ phenotype. Similarly, k_1_ and k_2_ values are lower for PICs assembled with G31R on UUG start codons than on AUG codons (k_1_ values 18 and 7 ×10^−3^ s^−1^, and k_2_ values 3.0 and 0.5 ×10^−3^ s^−1^, for AUG and UUG, respectively; Table 4A, rows 3-4 *versus* 1-2). The R_bound_ values also invert, becoming 0.19 and 0.20 for AUG and UUG, respectively (Table 4A, rows 3-4 *versus* 1-2). These results are consistent with the placement of the G31 residue directly across from the first position of the codon:anticodon helix (Figure 2A), where an arginine substitution could stabilize the formation of a U:U mismatch and the closed/P_IN_ state of the complex.

The additional Sui^−^ substitutions in the eIF5-NTD generated here, E26K, G29R, and N30R, all produced a similar pattern to what we observed for eIF5-G31R (Figures 4A-D). All three substitutions led to higher K_amp_ values in PICs with mRNA(UUG) versus mRNA(AUG) (Table 4A, rows 5-10), implying that these substitutions in eIF5, like G31R, shift the equilibrium more towards the closed/P_IN_ state at the near-cognate UUG start codon and away from it at AUG codons. As with eIF5-G31R, k_1_ and k_2_ were both lower on UUG codons than AUG codons for all three mutants (Table 4A). R_bound_ values were either equal for UUG and AUG codons (G29R) or greater for UUG codons (N30R and E26K), as was also the case for G31R (Table 4A). These results are consistent with the proposal that these positive charge substitutions in eIF5-NTD in the vicinity of the codon:anticodon helix electrostatically stabilize the P_IN_ conformation on near-cognate codons. This does not seem to be the only effect, however, because these substitutions actually appear to destabilize the closed/P_IN_ state on AUG codons, possibly due to steric clashes with the A:U base pair introduced by the arginine side chain.

We also determined the effects of substitutions in the eIF5-NTD at the same or nearby residues designed to decrease the positive charge or increase the negative charge: G29E, N30E, R28E and R28A. Because these substitutions produced hyperaccurate Ssu^−^ phenotypes in the genetic experiments described above, we examined their effects on eIF1A dissociation in the context of PICs assembled with eIF2 harboring the *SUI3-2* variant of the β-subunit (eIF2β-S264Y). Consistent with its Sui^−^ phenotype, we observed that *SUI3-2* eIF2 slows the rate of eIF1A dissociation from 48S PICs assembled on mRNA(UUG), reducing the differences in rate constants, K_amp_ and R_bound_ values between 48S PICs on AUG and UUG mRNAs, relative to the WT complexes (Figure 5A red circles *versus* red squares; row 2 Table 4A *versus* row 2 Table 4B). These results suggest that the *SUI3-2* mutant of eIF2 stabilizes the closed/P_IN_ state of the PIC at UUG codons.

**Figure 5.**
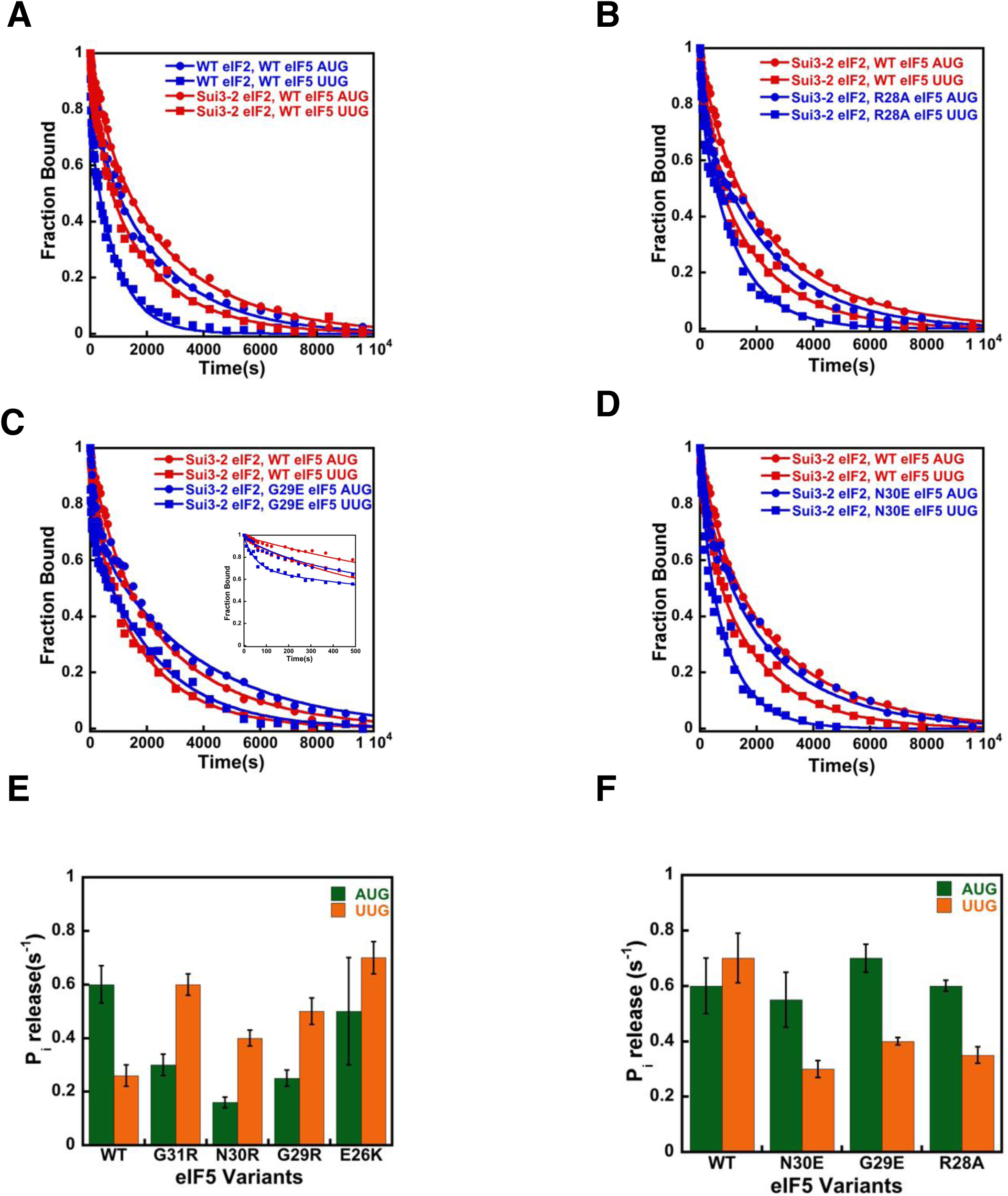
eIF5-NTD Ssu^−^ mutants destabilize the closed conformation of the PIC and accelerate P_i_ release at UUG codons in the presence of the *SUI3-2* Sui^−^ variant of eIF2. (A-D) eIF1A dissociation kinetics experiments conducted as in Figure 4 for PICs assembled with mRNAs containing AUG (closed circles) or UUG (closed squares) start codons and the following forms of eIF2 and eIF5: (A) WT eIF2/WT eIF5 (blue) or Sui3-2 eIF2/WT eIF5 (red); (B) Sui3-2 eIF2/WT eIF5 (red) or Sui3-2 eIF2/eIF5-R28A (blue); (C) Sui3-2 eIF2/WT eIF5 (red) or Sui3-2 eIF2/eIF5-G29E (blue); (D) Sui3-2 eIF2/WT eIF5 (red) or Sui3-2 eIF2/eIF5-N30E (blue). (E-F) Rates of P_i_ release from PICs assembled with mRNAs containing AUG (green) or UUG (orange) start codons and the indicated WT or mutant variants of eIF5. WT eIF2 was employed in (E), whereas Sui3-2 eIF2 was used in (F). All experiments were performed at least 2 times and error bars are calculated as average deviation.

With 48S PICs assembled with *SUI3-2* eIF2, the eIF5 variants R28A, G29E, and N30E increased the overall rate of eIF1A dissociation with mRNA(UUG) as compared to the native eIF5 (Figure 5B-D, blue squares versus red squares). These substitutions decrease the occupancy of the closed complex at UUG start codons, as indicated by the decreased K_amp_ values on UUG relative to AUG start codons compared to the case with *SUI3-2* PICs containing WT eIF5 (Table 4B, rows 5-10). The N30E and R28E derivatives of eIF5 also increase k_2_ values for complexes with UUG by ~2-fold (Table 4B, rows 3-4 and rows 7-10), indicating that these substitutions destabilize eIF1A binding to the closed state of the PIC at UUG start codons. All four Ssu^−^ eIF5 variants also increase k_1_ with both UUG and AUG codons, suggesting they destabilize eIF1A binding to the open state of the PIC. In all cases, the substitutions partially or completely restore the difference in R_bound_ values between AUG and UUG complexes that was eliminated by the *SUI3-2* mutant with WT eIF5 (Table 4B, rows 3-10 versus 1-2). Taken together, these results suggest that the Ssu^−^ eIF5 suppressors of *SUI3-2* eIF2 revert the equilibrium back towards the open/P_OUT_ conformation of the PIC at UUG codons while promoting the closed/P_IN_ conformation at AUG codons.

### eIF5-NTD substitutions at the codon:anticodon interface alter the coupling of P_i_ release to start codon recognition

We next checked the effect of the eIF5 substitutions on the rate of phosphate (P_i_) release from eIF2 in the PIC in response to recognition of cognate AUG and near-cognate UUG start codons. P_i_ release is a late step in start codon recognition and is gated by eIF1 release and movement of the eIF1A-CTT closer to the eIF5-NTD. It is thought to help commit the PIC to initiation at the selected point on the mRNA. Previous studies have shown that P_i_ release is influenced by the nature of the start codon in the mRNA, with a higher rate observed from PICs assembled on AUG start codons as compared to PICs on near-cognate UUG codons (Algire et al., 2005) (Saini et al., 2014).

In accordance with earlier studies (Saini et al., 2014; Algire et al., 2005), we observed that the kinetics of P_i_ release is 2-3-fold faster in response to AUG as compared to UUG start codons (k_obs_ values of 0.60 s^−1^ *versus* 0.26 s^−1^) (Figure 5E; Table 5A, row 1). This trend is reversed when the Sui^−^ G31R eIF5 mutant replaces the WT factor, with a k_obs_ of 0.6 s^−1^ for PICs assembled on UUG codons versus 0.3 s^−1^ for complexes on AUG codons (Figure 5E; Table 5A, row 2). This result is consistent with previous observations (Saini et al., 2014) and the Sui^−^ phenotype of the G31R mutant. Similarly, all of the new Sui^−^ substitutions in eIF5 (N30R, G29R, and E26K) suppress the rate of P_i_ release from the PIC in response to recognition of cognate AUG start codons and/or enhance it in response to near-cognate UUG codons (Figure 5E, compare green and orange bars; Table 5A, rows 3-5). These results are consistent with the conclusion that these substitutions that increase the positive charge on the eIF5-NTD in the region of the codon:anticodon helix stabilize the closed state of the PIC at UUG codons, while destabilizing it at AUG codons. Thus, in accordance with the eIF1A dissociation kinetics results described above, the P_i_ release kinetics for the positive charge eIF5 substitutions help explain their Sui^−^ phenotypes observed *in vivo*.

Next, we monitored the kinetics of AUG-and UUG-triggered P_i_ release from PICs assembled with the Sui^−^ *SUI3-2* eIF2 mutant and either WT eIF5 or one of the Ssu^−^ variants described above (N30E, G29E, R28A). In agreement with the eIF1A dissociation kinetics results described above, the *SUI3-2* mutant normalizes the rate of P_i_ release from complexes assembled on UUG and AUG start codons (k_obs_ values of 0.7 and 0.6 s^−1^, respectively; Figure 5F; Table 5B, row 1 *versus* Table 5A, row 1). Unlike the behavior of G31R eIF5, the *SUI3-2* mutant does not decrease the rate of P_i_ release with AUG start codons but instead only increases the rate with UUG codons (Figure 5F *versus* Figure 5E), suggesting that it specifically enhances the stability of the closed/P_IN_ conformation of the PIC at near-cognate (UUG) codons. Consistent with their effects in the eIF1A dissociation assay, the Ssu^−^ eIF5 variants N30E, G29E and R28A all suppress the effect of *SUI3-2* eIF2 by decreasing the rate of P_i_ release from UUG start codons ~2-fold (k_obs_ values between 0.3–0.4 s^−1^, Table 5B, rows 2-4) versus the rate observed with WT eIF5 (k_obs_ of 0.7 s^−1^, Table 5B, row 1), restoring the preference for AUG start codons. These results support the proposal that these substitutions, which increase the negative charge or decrease the positive charge in this region of the eIF5-NTD, destabilize the closed/P_IN_ state of the PIC at near-cognate codons.

### Stabilization of codon-anticodon interaction by the ribosome, eIF5, eIF2α and eIF1A

In contrast to the high conformational heterogeneity observed in previous reported py48S datasets (Llácer et al., 2015) (Hussain et al., 2014), the py48S-eIF5N is locked into a single configuration with the exception of small movements of eIF2 subunits γ, β, and domain 3 (D3) of the α subunit around the acceptor arm of the tRNA_i_ (see maps C1 and C2; Figures 6A,B and Figure 1 – figure supplement 1), and also of the flexible eIF3 domains at the solvent side of the 40S. While in our two previous studies (Llácer et al., 2015) (Hussain et al., 2014) the proportion of PIC complexes containing bound TC ranged from 1 to 15%, here it was around 40%. Although the reasons for the improvement are not entirely clear, in the study here, an optimal Kozak sequence was used in the mRNA and the complex was formed by pre-binding eIF4F to the mRNA. In any case, the improved conformational homogeneity and rigidity of the py48S is likely responsible for a significantly improved resolution in this study.

**Figure 6.**
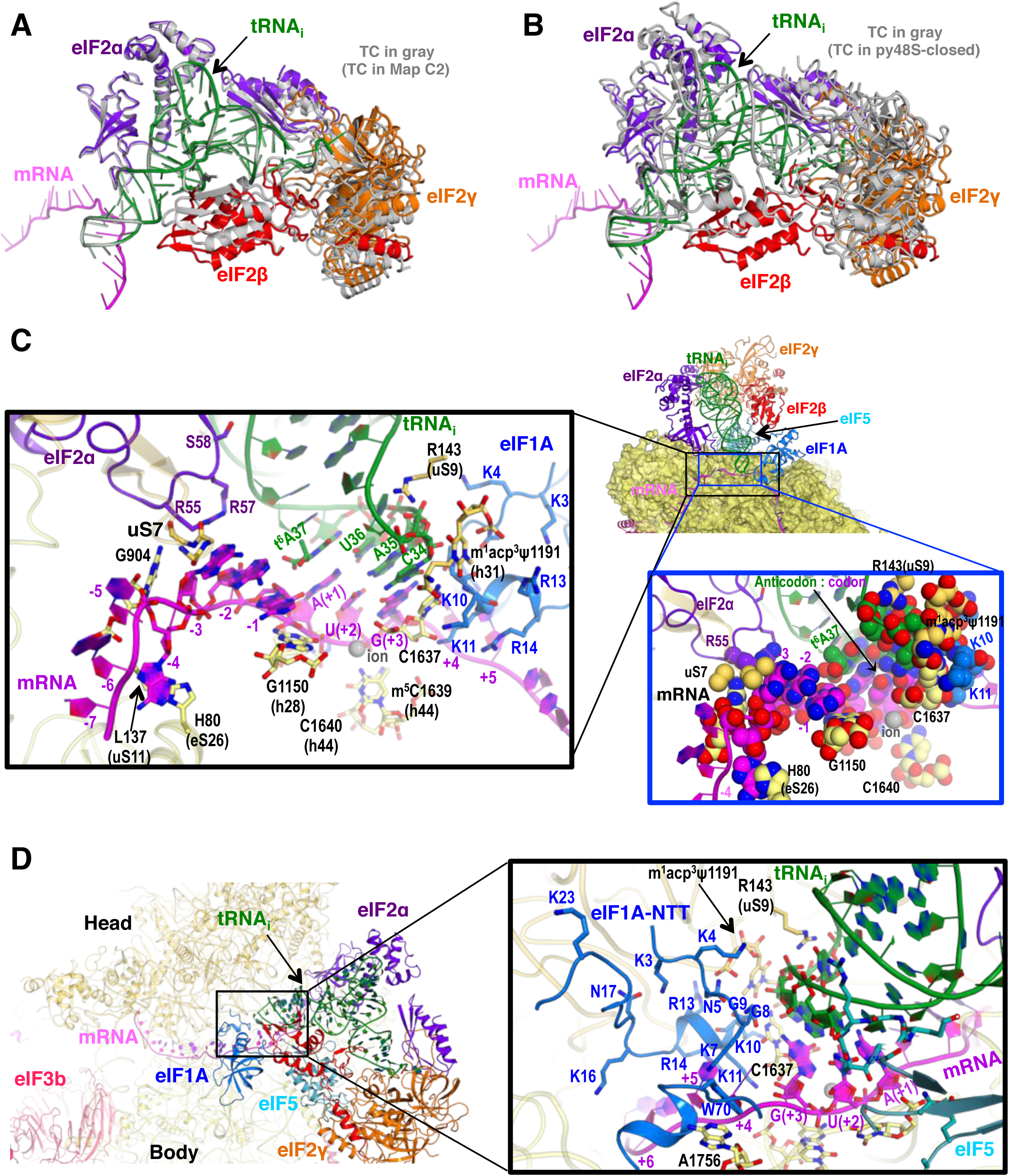
P-site conformation and surrounding elements in py48S-eIF5N. (A) Cartoon representation of the TC in Maps C1 and C2, resulting from superposition of the 40S body in the two maps. The tRNA_i_, mRNA, and each component of TC is colored differently for C1, whereas all components of C2 are in gray. (B) Cartoon representation of the TC in Map C1 and py48S-closed (PDB 3JAP), resulting from superposition of the 40S body in the two maps. The tRNA_i_, mRNA, and each component of TC is colored differently for C1, whereas all components from py48S-closed are in gray. (C) Cross-section of the 40S subunit along the mRNA path and tRNA_i_ bound in the P-site of py48S-eIF5N, viewed from the top of the 40S subunit. eIF1A, eIF2 and eIF5-NTD are also shown. Black box inset: Detailed view of the codon-anticodon and surrounding elements that stabilize this interaction. Ribosomal, tRNA_i_ and eIF2α residues involved in the interaction with mRNA at (minus) positions 5’ of the AUG codon (+1) are also shown. Blue box inset: Spheres representation of the same region in (C) highlighting the close packing of mRNA from positions -4 to +3 with its stabilizing residues of the ribosome, tRNA_i_ and eIF2α in the mRNA channel. (D) eIF1A NTT interactions with the codon-anticodon duplex and the 40S subunit in py48S-eIF5N.

In py48S-eIF5N, we observe an apparently greater accommodation of the codon-anticodon duplex in the P-site (Figure 2C) in concert with stabilization of the ASL by other elements from the eIF5-NTD (noted above), the 40S body, the eIF1A-NTT and eIF2α-D3 (Figure 6C). In this regard, the ribosomal elements (R143 of 40S protein uS9 and rRNA bases from h44 and h31) involved in the stabilization of the codon-anticodon duplex at the P-site described earlier for the py48S in the presence of eIF1 (Hussain et al., 2014) are also involved in py48S-eIF5N (Figure 6C). Moreover, a highly modified rRNA base at the 40S head (m_1_acp^3^ψ1191), not present in bacteria, plays a key role by providing a chair-like structure where the tip of the tRNA_i_ ASL (C34) sits. Mutations in an enzyme involved in modifying this base are associated with a severe syndrome in humans (Meyer et al., 2011). Also, an ion (revealed as a spherical density) (Figure 1 – figure supplement 3D) interacts with the phosphates of the U(+2) and G(+3) nucleotides of the AUG codon as well as the nearby rRNA residues G1150 and C1637, thereby playing a key structural role at the P-site (Figure 6C). As in previous py48S structures in the P_IN_ state (Llácer et al., 2015) (Hussain et al., 2014), the eIF1A-NTT interacts with the codon-anticodon helix via Gly8-Gly9, which also allow the NTT to loop back towards the P site (Figure 6D). We are now able to visualize the entire NTT unambiguously except for the terminal Met1 residue. The NTT occupies the cleft in between the head and body of the 40S around the A and P-sites, and several of its basic residues (Arg and Lys) establish contacts with rRNA residues from the 40S head and body (Figure 6D), essentially gluing them together to stabilize the closed conformation of the 48S. Recently, we established that substituting the conserved basic residues, as well as the yeast equivalents of eIF1A NTT residues identified as recurring substitutions in certain human uveal melanomas, decreases initiation at UUGs *in vivo* and selectively destabilizes PICs reconstituted at UUG codons *in vitro* (Martin-Marcos et al., 2017), as described above for eIF5-NTD Ssu^−^ substitutions.

### Interactions with the Kozak sequence of mRNA

In addition to the start codon, the mRNA bases at -1 to -4 in the E-site corresponding to the Kozak consensus sequence (Kozak, 1986), are locked into a single conformation in py48S-eIF5N. Bases -1 to -4 adopt an unusual but stable conformation, in which the adenine base at -4 is flipped out towards the 40S body, and the next three adenines (-3 to -1) stack with one another and are sandwiched by the uS7 β-hairpin loop and G1150 (from h28, at the neck of the 40S) (Figure 6C). Moreover, Arg55 of eIF2α interacts with the A nucleotide at -3, as previously reported (Hussain et al., 2014) (Pisarev et al., 2006); and the t^6^A37 base adjoining the tRNA_i_ anticodon interacts with the A at -1 through its threonylcarbamyol modification and also stacks with the adenine base at the +1 position of the start codon (Figure 6C). Interestingly, the absence of t^6^A37 in yeast increases translation initiation at upstream non-AUG codons (Thiaville et al., 2016), and therefore plays a role in stringent selection of AUG as start codon. In *S. cerevisiae*, A nucleotides at the -4 to -1 positions are highly preferred, particularly the A at -3 (Zur and Tuller, 2013), and are known to promote AUG recognition (Martin-Marcos et al., 2011; Hinnebusch, 2017). Placement of this favorable mRNA sequence at the E-site and its attendant interactions with elements of the 40S, eIF2α, and tRNA_i_ might pause the ribosome during scanning and help position the downstream AUG codon at the P-site. Its indirect stabilization of the codon-anticodon duplex and, therefore, of the closed conformation of the 48S might also facilitate dissociation of eIF1 after AUG recognition and subsequent eIF5-NTD binding.

### The path of mRNA and interaction with eIF3a at the mRNA exit channel of py48S-eIF5N

We could model the mRNA in the py48S-eIF5N from positions -14 to +17, spanning the entire mRNA channel plus additional nucleotides protruding from the two channel openings on the solvent side of the 40S (Figure 7A). The last 4 nucleotides at the 3’ end and first 14 nucleotides at the 5’ end of the mRNA could not be modeled owing to lack of unambiguous high-resolution density. As previously reported (Hussain et al., 2014), we observe kinks between the A and P codons and P and E codons (Figure 6C). At the mRNA entry site (Figure 7B), the latch is closed (Figure 1 – figure supplement 5) and the mRNA interacts with both elements from the head (uS3) and body (uS5 and h16) of the 40S. We showed recently that two conserved Arg residues in yeast uS3 in proximity to the mRNA (R116-R117) stabilize PIC:mRNA interaction at the entry channel, augmenting this activity of eIF3, and are crucial for efficient initiation at UUG codons and AUG codons in poor Kozak context *in vivo* (Dong et al., 2017). At the very 3’ end, the mRNA does not protrude away from the ribosome but instead points upward and remains attached to the 40S head. Whether this reflects the limited length at the 3’ end of the mRNA used here or represents the true trajectory of the mRNA 3’ end remains to be determined. The fact that proteins like eIF3g and eIF4B, which can interact with both mRNA and proteins of the 40S head (Cuchalova et al., 2010) (Walker et al., 2013), bind in this region may favor the latter hypothesis.

**Figure 7.**
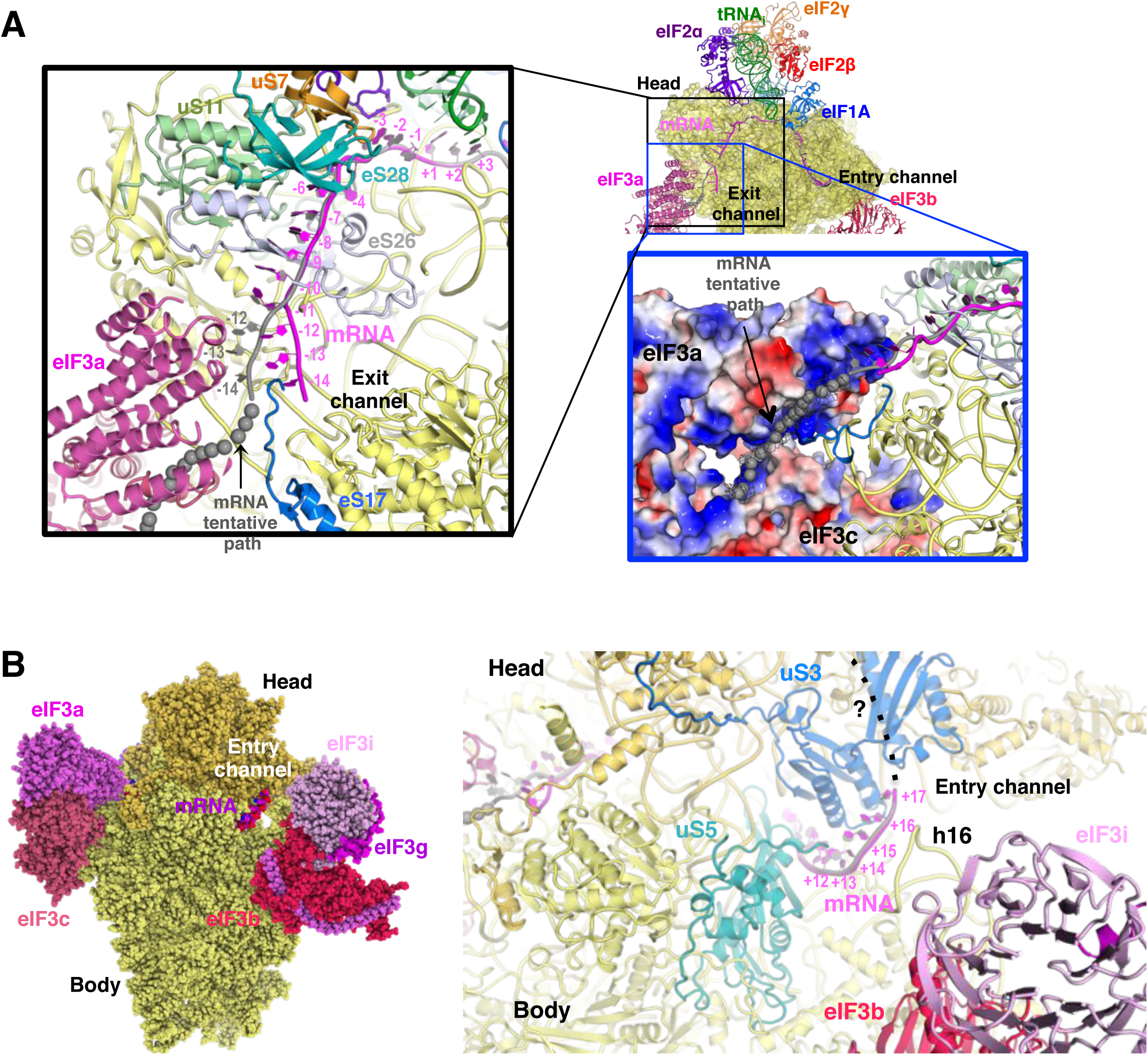
mRNA path at the exit and entry of the 40S mRNA channel in py48S-eIF5N. (A) Cross-section of the 40S subunit along the mRNA path of py48S-eIF5N, viewed from the top of the 40S subunit, showing both the entry and exit openings of the 40S mRNA channel. Path of the mRNA when the eIF3a/c PCI domains are present in high occupancy (ie. in model A) is shown in gray. Black box inset: mRNA at the exit tunnel (from nucleotides -1 to -13). mRNA interacting proteins uS7, uS11, eS17, eS26 and eS28 are colored orange, light green, blue, blue-white and teal, respectively. Grey spheres highlight a tentative path for the mRNA based on an unassigned density in Map A (see tubular-shaped density shown as a grey mesh in the blue box inset). Blue box inset: Surface electrostatic potential of eIF3a/c PCI domains (blue: basic; red: acidic) supports their proposed interaction with mRNA upstream of position -11 depicted as the tubular-shaped density shown in grey mesh. (B) mRNA at the entry tunnel opening (from nucleotides +11 to +17), colored as in A. mRNA-interacting 40S proteins uS3 and uS5 are colored blue and cyan, respectively. rRNA helix h16 also interacts with mRNA and is labeled. A possible trajectory for the 3’ end of the mRNA is proposed and shown as a discontinuous black line.

At the mRNA exit channel (Figure 7A), the mRNA interacts with uS7, us11, es17, eS26, eS28 and the 3’ end of the rRNA. At the exit channel pore, the -12 to -14 nucleotides of mRNA interact with the eIF3a-PCI domain, and interestingly this interaction seems to change the trajectory of the mRNA at its 5’end (Figure 7A, black inset), as seen in Map A (Figure 1 – figure supplement 1) containing higher eIF3 occupancy. In fact, an unassigned tubular density at the 5’ end of the modeled mRNA (Figure 7A, blue inset), which may correspond to part of the unmodeled first 14 nucleotides of the mRNA, approaches and lays on the electropositive surface of eIF3a. This possible interaction would be consistent with previous experiments showing cross-linking of mammalian eIF3a to the -11 to -17 positions in mRNA (Pisarev et al., 2008), and that the eIF3a-PCI is critical for stabilizing mRNA binding at the exit channel (Aitken et al., 2016).

### eIF3 architecture in the py48S-eIF5N complex

Only about 10% of the total number of particles have densities corresponding to various subunits of eIF3 (Figure 1 – figure supplement 1, Maps A and B). However, this proportion is higher than in any previous py48S complexes we have studied (Llácer et al., 2015) (Hussain et al., 2014). Except for the N-terminal helical bundle of eIF3c, which is still located at the subunit interface (Llácer et al., 2015) (Figure 8A), the eIF3 subunits in py48S-eIF5N are located entirely at the 40S solvent side, as in mammalian 43S structures and the yeast 40S-eIF1-eIF1A-eIF3 complex (des Georges et al., 2015) (Hashem et al., 2013) (Aylett et al., 2015). As in these previous structures, the eIF3a-eIF3c PCI heterodimer sits near the mRNA exit tunnel; whereas the quaternary complex of eIF3b/eIF3i/eIF3g/eIF3a-Cter is found near the mRNA entry channel, where the eIF3b WD40 β-propeller contacts uS4, h21, eS6c and h16 (Figure 8A, B), slightly altering h16 conformation (Figure 8E). Although these eIF3 submodules are roughly in the same locations in py48S-eIF5N, we observe some differences in comparison to previous structures (Aylett and Ban, 2017; Valášek et al., 2017). In py48S-eIF5N, the eIF3a-eIF3c PCI domain is displaced laterally, bringing more of the PCI domain closer to the 40S. This altered conformation is probably due to its interaction with mRNA (Figure 8 A,D). The position of eIF3i and the eIF3b-RRM domain with respect to the eIF3b β-propeller is also slightly different (Figure 8C), probably reflecting the connections of these subdomains to the eIF3b β-propeller by flexible linkers. The eIF3a-Cterm helix likely also helps to position the different eIF3b domains because the beginning 2/3^rd^ of the helix runs underneath the eIF3b β-propeller and the last 1/3rd interacts with the β-sheet face of the eIF3b-RRM domain (Figure 8B). The local resolution of eIF3 at this location prevents unambiguous assignment of residues to the eIF3a-Cterm helix.

**Figure 8.**
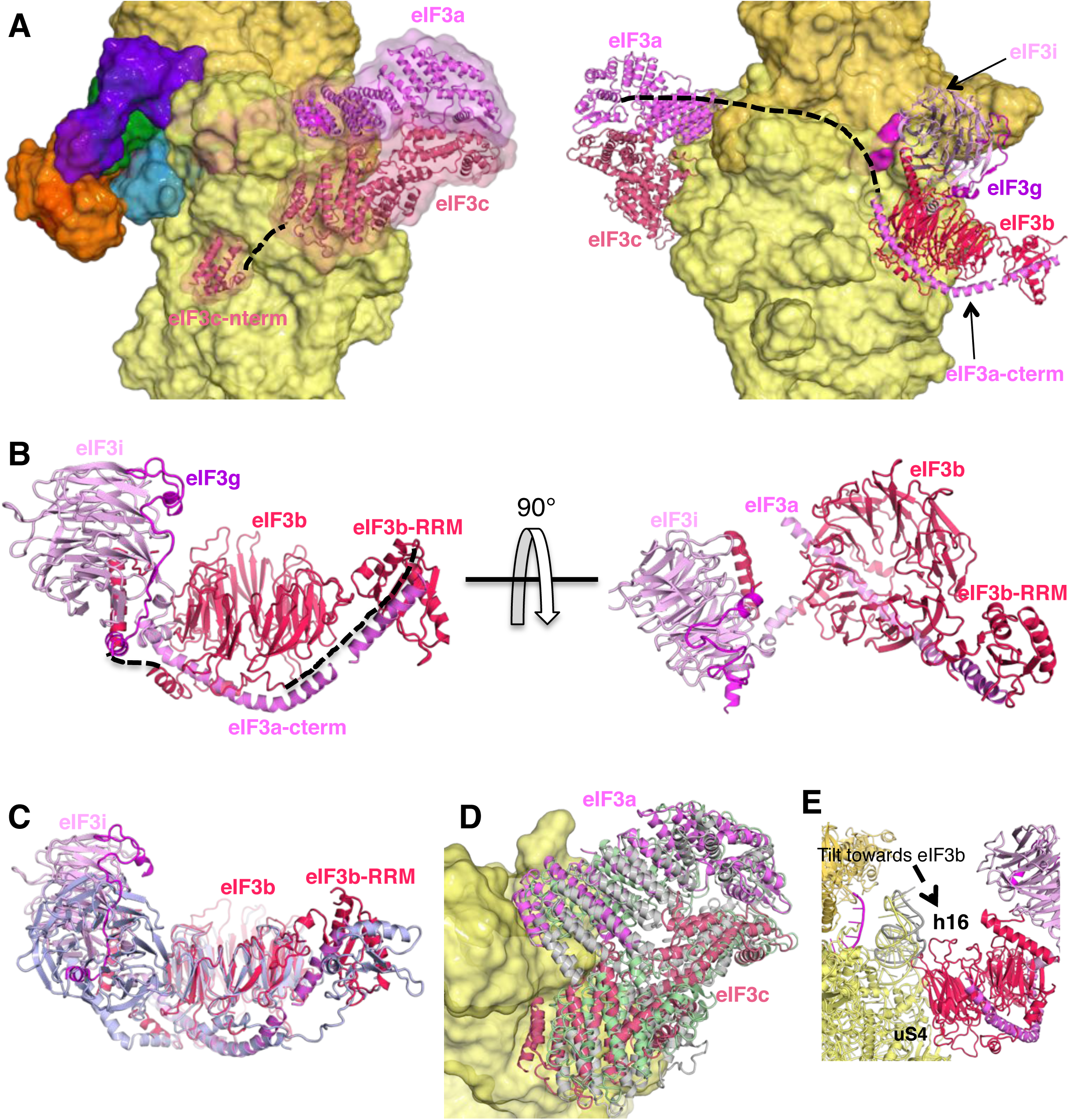
eIF3 architecture within py48S-eIF5N. (A) Two different views of the py48S-eIF5N PIC showing the locations of the different eIF3 subunits. All eIF3 domains, except for the eIF3c N-terminal helical bundle, reside on the solvent-exposed side of the 40S subunit. Shown as dashed black lines are the proposed linker connecting the eIF3c helical bundle and PCI domain; and the proposed path for the central part of eIF3a connecting the eIF3a/PCI domain and eIF3a C-term helix, of which the latter interacts with the eIF3b β-propeller. (B) Quaternary complex eIF3b/eIF3i/eIF3g/eIF3a-Cterm, shown in two different orientations. Unresolved connections between the eIF3b β-propeller and the eIF3b C-terminal helix and eIF3b RRM domain are shown as dashed black lines. (C) Superimposition of the quaternary complex in (B) to its counterpart in the mammalian 43S complex (light blue; PDB 5A5U), by aligning the two eIF3b β-propeller domains, shows a slightly different arrangement of the eIF3b RRM and eIF3i β-propeller with respect to the eIF3b β-propeller in the two complexes. (D) Cartoon representation of a superimposition of the eIF3a/c PCI domains in the py48S-eIF5N (Map A), py48S-closed (green; PDB 3JAP), and yeast 40S-eIF1-eIF1A-eIF3 complex (grey; PDB 4UER) resulting from aligning the 40S bodies. (E) Superimposition of py48S-eIF5N from Maps 1 and B, aligned according to 40S body, showing a displacement of h16 towards the eIF3b β-propeller in Map B (shown in gray).

## Discussion/Conclusion

In previous structures of yeast PICs containing Met-tRNA_i_ base-paired to an AUG codon, the gatekeeper molecule eIF1 is still bound to the 40S platform, indicating that these structures likely depict intermediate states in the pathway prior to P_i_ release from eIF2-GDP-P_i_, which is gated by eIF1 dissociation. In the py48S and py48S-closed structures, where tRNA_i_ is tightly enclosed in the P site, the location and conformation of the β-hairpin loops 1 and 2 of eIF1 are different from their counterparts in both the py48S-open complex and the simpler PIC containing only eIFs 1 and 1A. These changes to eIF1 are required to accommodate tRNA_i_ binding in the conformation observed in the previously reported py48S closed complexes (Llácer et al., 2015) (Hussain et al., 2014). The remodeling of eIF1 loop-1 disrupts certain interactions anchoring eIF1 to the 40S platform, presumably as a prelude to its eventual dissociation. Here, we describe a PIC representing a step further in the pathway, in which eIF1 has dissociated and is replaced by the eIF5-NTD on the 40S platform (Movie 1). The eIF5-NTD interacts directly with the codon:anticodon duplex and might act to stabilize selection of an AUG start codon, as well as allowing greater accommodation of Met-tRNA_i_ in the P-site in a tilted conformation. The altered location and conformation of Met-tRNA_i_ in the py48S-eIF5N complex is completely incompatible with eIF1 binding to the platform, even in the distorted state observed in the previous closed complexes. The eIF5-NTD, in contrast, is complementary to this new conformation and makes stabilizing interactions with the tRNA_i_, thus promoting the fully closed/P_IN_ state of the PIC and preventing rebinding of eIF1. The position and tilted conformation of Met-tRNA_i_ in the py48S-eIF5N complex appears to set the stage for the next step in the initiation pathway, eIF5B-catalyzed subunit joining.

As shown previously (Conte et al., 2006), eIF5-NTD and eIF1 share the same protein fold, and they mostly contact the 40S subunit using analogous structural elements. Moreover, eIF1 and the structurally similar portion of eIF5-NTD contact essentially the same surface of the 40S platform. However, whereas eIF5-NTD binding is compatible with the more highly accommodated P_IN_ state of Met-tRNA_i_ and its tilting towards the 40S body observed in the py48S-eIF5N structure – and, in fact, appears to stabilize this fully accommodated state -eIF1 would clash extensively with Met-tRNA_i_ in this location and orientation. The β-hairpin loops 1 and 2 of eIF5-NTD make extensive, favorable contacts with the Met-tRNA_i_, and because eIF5 loop-2 is shorter/more basic and oriented away from the Met-tRNA_i_, it avoids the electrostatic clash with the tRNA_i_ D-loop predicted for the larger/more acidic loop-2 of eIF1 that projects towards the tRNA_i_ (Figure 2C). Supporting this view, we recently demonstrated that substituting acidic and bulky hydrophobic residues in eIF1 loop-2 with alanines or basic residues increases UUG initiation in vivo and stabilizes TC binding to the PIC at UUG codons, as would be expected from eliminating electrostatic/steric clashing, or introducing electrostatic attraction, between eIF1 loop-2 and tRNA_i_, which removes an impediment to the P_IN_ state to enhance selection of a near-cognate start codon (Thakur and Hinnebusch, 2018).

Our genetic findings provided evidence that electrostatic contacts of basic residues R28 and R73 in eIF5-NTD loops 1 and 2, respectively, with different segments of the tRNA_i_ ASL stabilize the closed/P_IN_ conformation at the start codon, as replacing them with Ala or Glu residues reduced initiation at UUG codons in yeast cells harboring the hypoaccurate Sui^−^ variant eIF2β-S264Y. Consistent with this, the eIF5-R28A substitution also disfavored and destabilized the closed PIC conformation as judged by eIF1A dissociation kinetics, and decreased P_i_ release from eIF2-GDP-P_i_, primarily or exclusively at UUG codons in 48S PICs reconstituted with the eIF2β-S264Y variant. Similar findings were observed on introducing acidic residues at G29 and N30 in the eIF5-NTD loop-2. The relatively greater effects of these substitutions on UUG initiation *in vivo*, and in disfavoring/destabilizing the closed PIC conformation and reducing the rate of P_i_ release *in vitro* at UUG versus AUG codons, taken in combination with earlier findings that the closed/P_IN_ state of the PIC is inherently less stable at UUG versus AUG codons (Maag et al., 2006); Kolitz et al 2009), suggests that non-canonical 48S PICs at UUG codons are more sensitive to loop-2 mutations that disrupt electrostatic stabilization of the codon:anticodon duplex by the eIF5-NTD loop-2. Thus, our structural, genetic and biochemical evidence all suggest that contacts between the eIF5-NTD and the tRNA_i_ ASL promote the closed/P_IN_ conformation of the PIC, and thus are particularly important for efficient initiation not only at AUG codons but also at a near-cognate UUG codon, whereas eIF1 binding in virtually the same location on the 40S platform destabilizes the closed/P_IN_ state and enforces a requirement for the perfect codon:anticodon duplex formed at AUG codons.

We found that introducing basic Lys or Arg residues at G29, N30, and E26, which should increase electrostatic attraction between eIF5-NTD loop-2 and the codon:anticodon helix, had the opposite effects of acidic substitutions at G29 and N30 at UUG codons, increasing the UUG:AUG initiation ratio *in vivo*, and favoring/stabilizing the closed PIC conformation and increasing the rate of P_i_ release at UUG codons *in vitro*. These phenotypes mimicked those of the G31R substitution (Maag et al., 2006)—the first described Sui^−^ substitution in eIF5 encoded by *SUI5*. These results seem to indicate that increasing electrostatic attraction with loop-2 can stabilize mismatched codon:anticodon duplexes and thereby promote initiation at near-cognate start codons. However, G31R does not increase initiation at other near-cognate triplets besides UUG *in vivo* (Huang et al., 1997), and it disfavors the closed PIC conformation and slows down P_i_ release at AUG codons *in vitro* (Saini et al., 2014) (Figures 4 and 5). Thus, the putative electrostatic stabilization seems to apply exclusively to the mismatched UUG:anticodon duplex and not those formed by other near-cognates, and to have the opposite effect on the perfect codon:anticodon helix formed at AUG. More work is required to understand the molecular basis of this exquisite specificity of basic loop-2 substitutions for UUG codons as well as the relative specificity for other non-canonical start codons.

The affinity of eIF1 for the free 40S subunit is ~30-fold higher than that of the eIF5-NTD (Cheung et al., 2007) (Nanda et al., 2013), which would favor initial binding of eIF1 over eIF5. However, the affinity of eIF1 for the 43S·mRNA(AUG) complex is ~20-fold lower (Cheung et al., 2007), whereas the affinity of eIF5 for the same complex is >300-fold higher, in comparison to their respective affinities for the free 40S subunit (Algire et al., 2005). As a result, following AUG recognition, the affinity of eIF5 for the PIC exceeds that of eIF1 by two orders of magnitude. The relatively high rate of eIF1 dissociation from the PIC on AUG recognition (Maag et al., 2005) should allow the eIF5-NTD to compete with eIF1 for re-binding to the 40S platform, and the relatively higher affinity of eIF5 for the 43S·mRNA(AUG) complex should favor the replacement of eIF1 by eIF5-NTD on the platform.

The notion that the eIF5-NTD and eIF1 compete for binding to the PIC helps to explain previous findings in yeast (Valasek et al., 2004; Nanda et al., 2009; Martin-Marcos et al., 2011) and mammals (Ivanov et al., 2010) (Loughran et al., 2012; Terenin et al., 2016) that overexpressing eIF1 or eIF5 have opposing effects on initiation accuracy, with overexpressed eIF1 increasing discrimination against near-cognate triplets or AUGs with sub-optimal Kozak sequences, and overexpressed eIF5 boosting utilization of poor start codons. This effect of overexpressed eIF1 implies that eIF1 dissociation from the PIC does not necessarily lead to an immediate release of P_i_, an irreversible reaction; a fast rate of eIF1 re-binding driven by mass action can allow resumption of scanning and prevent P_i_ release, consistent with the faster rate of eIF1 release than P_i_ release from WT PICs reported previously (Nanda et al., 2013). By the same logic, overexpressed eIF5 would decrease reassociation of eIF1 through increased competition with the eIF5-NTD for binding to the 40S platform, and thus stimulate P_i_ release at poor initiation sites. In addition to increasing competition by the eIF5-NTD with eIF1 for 40S-binding, eIF5 overexpression might also enhance the ability of the eIF5-CTD to more actively evict eIF1 from the 40S by competing for binding to the beta subunit of eIF2 (Nanda et al., 2009; Llácer et al., 2015).

The inference that P_i_ release does not occur immediately on eIF1 dissociation is also supported by evidence of a functional interaction between the eIF5-NTD and eIF1A-CTT, involving movement of these domains towards one another in the PIC, which is required for rapid P_i_ release on AUG recognition following the dissociation of eIF1 from the 40S subunit (Nanda et al., 2013). We previously speculated that movement of the eIF5-NTD towards eIF1A on AUG recognition would involve replacement of eIF1 with eIF5-NTD on the platform, based on the structural similarity between eIF1 and the eIF5-NTD and also evidence that eIF1 is in proximity to the eIF1A-CTT in the open, scanning conformation of the PIC but moves away from eIF1A on AUG recognition (Maag et al., 2005) at essentially the same rate that the eIF5-NTD and eIF1A-CTT move towards one another in the PIC (Nanda et al., 2013). The presence of the eIF5-NTD on the platform observed here in the closed py48S-eIF5N complex, in essentially the same location occupied by eIF1 in the 40S·eIF1·eIF1A and py48S-open complexes (Llácer et al., 2015), provides strong structural evidence supporting our previous hypothetical model.

The GAP activity of eIF5 is dependent on Arg-15 at the N-terminal end of the NTD. Although Arg-15 is in the vicinity of the GDPCP bound to eIF2γ in py48S-eIF5N, it is too distant to function in stimulating GTP hydrolysis. Previous evidence indicates that GTP hydrolysis can occur in the scanning PIC and that P_i_ release rather than hydrolysis *per se* is the step of the reaction most highly stimulated by AUG recognition. The position of eIF5-NTD in our structure is compatible with the notion that GTP hydrolysis has already occurred by the time the eIF5-NTD replaces eIF1 on the platform because the eIF5-NTD is engaged with the codon:anticodon helix rather than the GTP-binding pocket of eIF2γ (Figure 2 – figure supplement 1D). Movement of the eIF5-NTD into the eIF1 binding site might be the trigger that releases P_i_ from eIF2γ and could be the movement detected previously that brings the eIF5-NTD closer to the eIF1A-CTT upon start codon recognition, a conformational change that is crucial for allowing P_i_ release (Nanda et al., 2013). The presence of eIF1 on the 40S platform of the scanning PIC would potentially block access of the eIF5-NTD to the GTP molecule bound to eIF2γ. However, Arg-15 resides within an extended NTT devoid of secondary structure that would likely be flexible enough to allow insertion of Arg-15 into the eIF2γ GTP-binding pocket when the eIF5-NTD is tethered to the PIC via known interactions of the eIF5-CTD with the eIF3c NTD or eIF2β NTT (Kirillov et al., 2002).

The more accommodated position of Met-tRNA_i_ and its tilting towards the 40S body observed with eIF5-NTD bound at the P-site in py48S-eIF5N appears to set the stage for the binding of eIF5B and attendant subunit joining, as this tRNA_i_ location/conformation is similar to that found in 80S initiation complexes with bound eIF5B (Figure 2 – figure supplement 2A) (Fernandez et al., 2013; Yamamoto et al., 2014). A recent study indicates that eIF5 and eIF5B cooperate in recognition of the start codon (Pisareva and Pisarev, 2014). The stabilizing effect of the eIF5-NTD on the more fully accommodated/tilted conformation of Met-tRNA_i_ in the P-site may form the basis of cooperation between eIF5 and eIF5B in recognition of AUG codons. In fact, the presence of the eIF5-NTD on the platform does not seem to impose any steric hindrance to the binding of eIF5B (Figure 2 – figure supplement 2B). Interestingly, the tRNA_i_ conformation in the py48S-eIF5N complex closely resembles that observed in bacterial translation initiation complexes containing IF2, the eIF5B homolog, suggesting that the processes of tRNA_i_ accommodation in the P-site converge on a similar position and orientation of tRNA_i_ in the final stages of eukaryotic and bacterial translation initiation (Figure 2 – figure supplement 2A,C,D). The mRNA path in the mRNA channel, mainly from -8 to +4 positions (Figure 2 – figure supplement 2F,G), is also surprisingly similar between bacterial and eukaryotic initiation complexes.

Both major subdomains of eIF3 are visible in py48S-eIF5N almost exclusively on the solvent side of the 40S, as previously observed in mammalian 43S structures and the yeast 40S-eIF1-eIF1A-eIF3 complex (des Georges et al., 2015) (Hashem et al., 2013) (Aylett et al., 2015). The eIF3a-eIF3c PCI domain is located at the mRNA exit tunnel and seen for the first time interacting with the -12 to -14 mRNA nucleotides, consistent with our recent biochemical analyses implicating the eIF3a-NTD in interaction with this region of the mRNA (Aitken et al., 2016). The quaternary complex of eIF3b/eIF3i/eIF3g/eIF3a-Cterm is located near the mRNA entry channel in the vicinity of the +12 to +17 mRNA nucleotides, which is also consistent with our recent biochemical studies. Comparison of py48S-eIF5N to the previous py48S-open and py48S-closed structures containing eIF1 (Llácer et al., 2015) reveals a dramatic alteration of the position of this eIF3 quaternary complex, which moves from the entry channel on the solvent side in py48S-eIF5N to communicate directly with initiation factors on the interface surface of the 40S subunit in py48S-open and py48S-closed complexes (Figure 9A-C). We originally interpreted the density map of py48S-closed as indicating binding of the eIF3i β-propeller in association with the 3g-NTD and 3b-CTD beneath the eIF2γ subunit of TC (Llácer et al., 2015). More recently it was suggested that the drum-like density attributed to the 3i β-propeller is actually the larger β-propeller of 3b, and that the density in contact with eIF1 is the 3b-RRM rather than the eIF3c-NTD (Simonetti et al., 2016). Based on a more recent, higher resolution map of a new py48S-open complex (unpublished observations, PDB 6GSM) we agree with these reassignments. The position of the eIF3b RRM at the interface surface of the py48S-open and -closed complexes, where it can contact eIF1, would clash with the eIF5-NTD bound to the platform in place of eIF1 in py48S-eIF5N (Figure 9D-E), which likely contributes to relocation of the eIF3b/eIF3i/eIF3g/eIF3a-Cterm module to the 40S solvent side, as observed here in the py48S-eIF5N complex.

**Figure 9.**
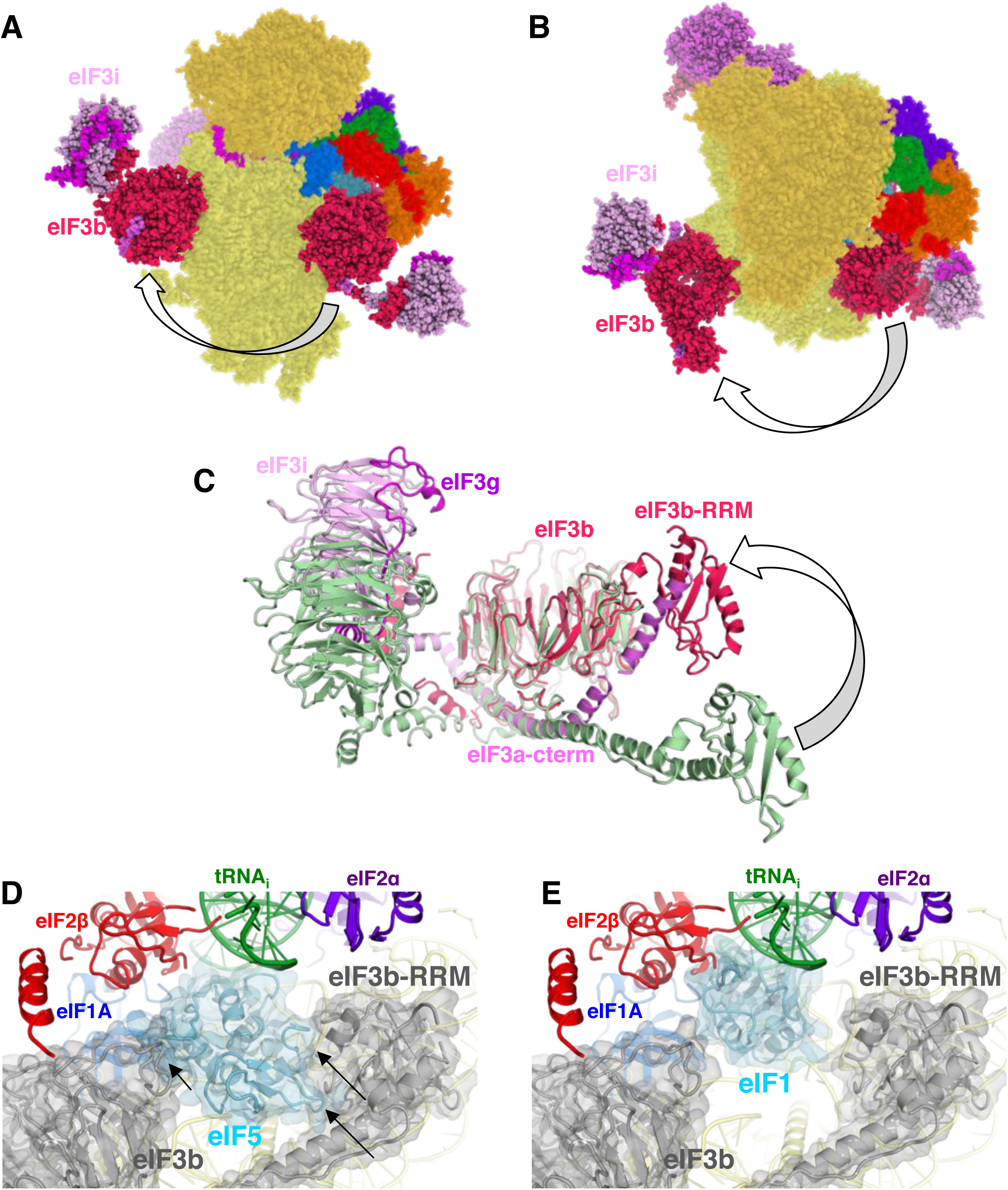
eIF3 conformational changes. (A,B) Side and top views of a composite representation showing the two different locations reported for the quaternary complex eIF3b/eIF3i/eIF3g/eIF3a-Cterm on the solvent or subunit surfaces of the 40S subunits in py48S-eIF5N (Map A or B) and an unpublished py48S-closed structure with eIF1 still present on the 40S platform in place of eIF5-NTD (PDB 6GSN). The arrow shows the direction of the rearrangement of this eIF3 subcomplex from the interface to solvent surface, which probably occurs after AUG recognition, complete accommodation of tRNA_i_ and eIF1 dissociation from the 48S. (C) Superimposition of the eIF3b/eIF3i/eIF3g/eIF3a-Cterm quaternary complex observed in py48S-eIF5N with that found on the 40S subunit interface in py48S-closed (in light green; PDB 6GSN), aligning the eIF3b β-propellers, shows how this eIF3 subcomplex undergoes internal rearrangements in the transition between the two states, possibly resulting from constraints imposed by its interactions with eIF2 and eIF1 unique to the subunit interface location. (D) Modeling of the eIF3b/eIF3i/eIF3g/eIF3a-Cterm quaternary complex observed in py48S-closed (PDB 6GSN) at the subunit interface of py48S-eIF5N. The location of eIF5-NTD in the latter complex seems to be incompatible with this position of eIF3b/eIF3i/eIF3g/eIF3a-Cterm at the subunit interface, clashing with it at multiple points (highlighted with black arrows). (E) The eIF3b/eIF3i/eIF3g/eIF3a-Cterm quaternary complex in the py48S-closed complex (PDB 6GSN) shows no clashes with eIF1 bound at the P site.

As noted above, the solvent-side location of the eIF3b/eIF3i/eIF3g/eIF3a-Cter module has also been observed in yeast and mammalian PICs lacking mRNA, and we proposed previously (Llácer et al., 2015) that the transition of this module from the solvent side to the interface surface might be triggered by PIC attachment to the mRNA, and that its interactions with initiation factors in the decoding center could help prevent dissociation of mRNA from the mRNA binding cleft during scanning. Thus, the view emerges that transition of the eIF3b/eIF3i/eIF3g/eIF3a-Cterm module from the solvent side to the interface side of the 40S occurs at the commencement of the scanning process and is reversed following AUG recognition and the replacement of eIF1 by the eIF5-NTD on the 40S platform.

The results of this study allow us to propose a model (Figure 10) of probable steps in translation initiation after the recognition of the start codon but prior to the subunit joining. Base pairing of Met-tRNA_i_ with the AUG start codon in the P_IN_ state of the closed conformation of the 48S PIC weakens eIF1 binding to the 40S subunit. Following eIF1 dissociation and attendant loss of its interaction with the eIF3b RRM, the eIF3b/eIF3i/eIF3g/eIF3a-Cter module relocates back to the solvent side of the 40S and allows binding of the eIF5-NTD at the site vacated by eIF1. The Met-tRNA_i_ binds more deeply in the P site and tilts towards the 40S body in a manner stabilized by extensive interactions of eIF5-NTD loop-1/loop-2 residues with the AUG:anticodon duplex. The eIF5-NTD also prevents re-binding of eIF1 and a return to the scanning conformation of the PIC. As proposed previously (Nanda et al., 2013), the eIF5-NTD functionally interacts with the eIF1A-CTT to permit irreversible P_i_ release and subsequent dissociation of eIF2-GDP. eIF5B is recruited via interaction with the extreme C-terminus of the eIF1A-CTT, captures the accommodated tRNA_i_ in the P-site and stimulates joining of the 60S subunit to produce the 80S initiation complex, leading to the final stage of initiation.

**Figure 10.**
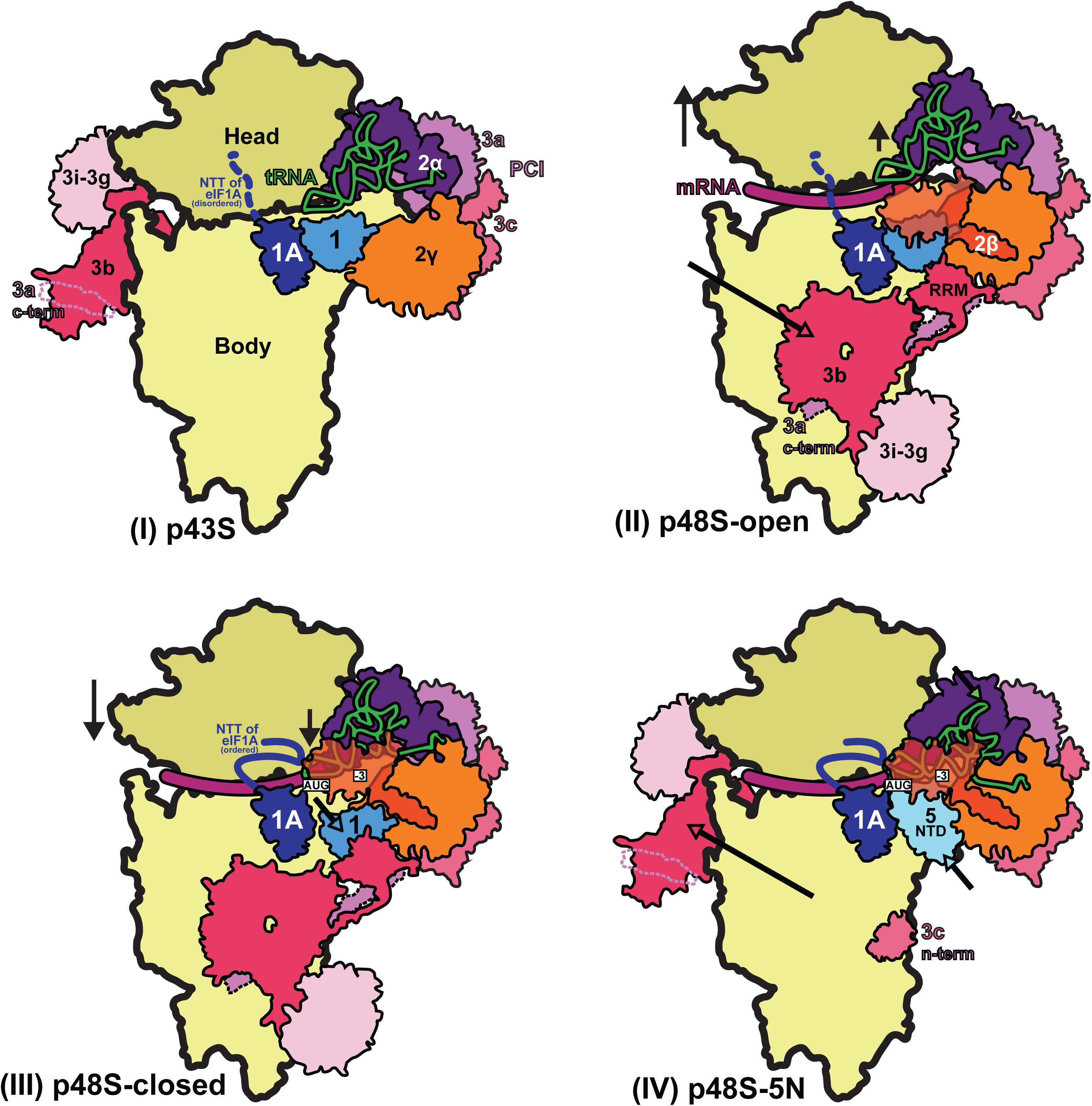
Schematic model of major conformational changes during initiation. (I) Binding of eIF1, eIF1A and eIF3 to the 40S subunit facilitates TC binding in the P_OUT_ conformation to form the 43S PIC. The disordered NTT of eIF1A is shown as a dashed line. eIF2β and 3c N-term were not resolved in this complex. (II) Upon mRNA recruitment facilitated by the upward movement of the 40S head (and also Met-tRNAi; shown by black arrows), which expands the mRNA entry channel, eIF3 undergoes major conformational changes and the eIF3b/eIF3i/eIF3g/eIF3a-Cterm module is repositioned from its initial position on the solvent exposed face of the 40S to the subunit-interface (shown by a pink arrow), contacting eIF2γ and eIF1. These contacts, together with eIF2β contacts with eIF1 and Met-tRNAi, probably stabilize this open scanning-competent conformation of the PIC (py48S-open). eIF2β is semitransparent overlapping eIF1.) (III) After recognition of the start codon, the 40S head moves downward (shown by a black arrow) to clamp in the mRNA and the ASL of the Met-tRNA_i_ goes deeper into the P site (P_IN_ state). eIF2β loses contact with eIF1 and Met-tRNAi and moves away to interact with the 40S head, the NTT of eIF1A stabilizes the codon:anticodon duplex, and eIF2α-D1 interacts with the -3 position of mRNA in the E site. eIF1 is also slightly displaced by Met-tRNA_i_ from its original position on the 40S platform (shown by a cyan arrow). (IV) Base pairing of Met-tRNA_i_ with the AUG start codon in the P_IN_ state and Met-tRNA_i_ tilting towards the 40S body (shown by a green arrow) weakens eIF1 binding to the 40S subunit causing eIF1 dissociation and, consequently, the eIF3b/eIF3i/eIF3g/eIF3a-Cter module relocates back to the solvent side of the 40S (shown by a pink arrow) and the eIF5-NTD binds at the site vacated by eIF1 (shown by a light cyan arrow) and stabilizes this Met-tRNA_i_ conformation through extensive interactions of eIF5-NTD loop-1/loop-2 residues with the AUG:anticodon duplex. In subsequent steps (not shown), eIF5B binds and captures the accommodated Met-tRNA_i_ in the P-site and stimulates the joining of the 60S subunit to produce the 80S initiation complex, leading to the final stage of initiation.

## Experimental Procedures

### Purification of Ribosomes, mRNA, tRNA, and Initiation Factors

*Kluyveromyces lactis* 40S subunits were prepared as described earlier (Fernandez et al., 2014). *Saccharomyces cerevisiae* eIF3 and eIF2 were expressed in yeast while eIF1, eIF1A, eIF5, eIF4A and eIF4B were expressed in *Escherichia coli* as recombinant proteins and purified as described (Acker et al., 2007) (Mitchell et al., 2010). eIF4G1 was co-expressed with eIF4E in *E. coli* and both purified as a complex as previously described (Mitchell et al., 2010) with modifications. A Prescission cleavage site was introduced just before the eIF4G1 coding sequence on the reported expression plasmid (Mitchell et al., 2010) in order to cleave and remove the GST tag from eIF4G1. Wild type tRNA_i_ was overexpressed and purified from yeast and aminoacylated as described (Acker et al., 2007). The mRNA expression construct comprised a T7 promoter followed by the 49-nt unstructured mRNA sequence of 5ʹ-GGG[CU]3[UC]4UAACUAUAAAA<u>AUG</u>[UC]_2_UUC[UC]4GAU-3ʹ (with start codon underlined), cloned between XhoI and NcoI sites in a pEX-A2 plasmid (Eurofins Genomics). AUG context optimality was inferred from the sequences of highly expressed genes in yeast, as reported in (Zur and Tuller, 2013). The mRNA was produced by T7 run-off transcription of the plasmid linearised by EcoRV (restriction site embedded in the mRNA sequence) according to a standard protocol (???REF). A 2mL transcription reaction was resolved by electrophoresis on an 8M Urea, 12% acrylamide gel. A single mRNA band, visualized by UV light, was excised from the gel and mRNA was electro-eluted in TBE buffer, concentrated and buffer exchanged by dialisis into storage buffer (10mM ammonium acetate, pH5.0, 50 mM KCl). mRNA was capped with Vaccinia Capping System (New England Biolabs, M2080S) according to the manufacturer’s instructions and further purified on an 8M Urea, 12% acrylamide gel as described above. The final concentration of mRNA was determined by A_2_60 measurement.

### Reconstitution of the 48S complex

First, a 43S mix was reconstituted by incubating 95 nM 40S subunits with eIF1, eIF1A, TC (consisting of eIF2, GDPCP and Met-tRNA_i_), eIF3 and eIF5 in 40S:eIF1:eIF1A:TC:eIF3:eIF5 molar ratios of 1:2.5:2.5:2:2:2.5, in 20 mM MES, pH 6.5, 80 mM potassium acetate, 10 mM ammonium acetate, 5-8mM magnesium acetate, 2mM dithiothreitol (DTT), 1 µM zinc acetate. Separately, an mRNA-eIF4 complex was prepared, containing eIF4G1, eIF4E, eIF4A, eIF4B and capped mRNA in molar ratios of 1.5:1.5:5:2:2.5 with respect to the 40S ribosome in the final 48S mix (see below), in 20 mM Hepes, pH 7.4, 100 mM potassium chloride, 5mM magnesium chloride, 2mM DTT, 3mM ATP). The volume of the mRNA-eIF4 mix was 5 times smaller than the 43S mix volume. Both the 43S mix and the mRNA-eIF4 mix were incubated separately for 5 min at room temperature before mixing them together to produce a 48S mix. After incubation for 2 min at room temperature, the sample (at a 40S final concentration of 80 nM) was cooled to 4°C and used immediately to make cryo-EM grids without further purification. When formaldehyde was used to crosslink the 48S complex (as described below), a solution at 3% in 48S mix buffer (at 1% final concentration of formaldehyde) was used just prior to making the cryo-EM grids.

### Electron microscopy

Three µl of the 48S complex was applied to glow-discharged Quantifoil R2/2 cryo-EM grids covered with continuous carbon (of ~50Å thick) at 4 °C and 100% ambient humidity. After 30s incubation, the grids were blotted for 2.5-3 s and vitrified in liquid ethane using a Vitrobot Mk3 (FEI).

Automated data acquisition was done using the EPU software (FEI) on a Tecnai F30 Polara G2 microscope operated at 300 kV under low-dose conditions (35 e^−^/Å_2_) using a defocus range of 1.2 – 3.2 µm. Images of 1.1 s/exposure and 34 movie frames were recorded on a Falcon III direct electron detector (FEI) at a calibrated magnification of 104,478 (yielding a pixel size of 1.34 Å). Micrographs that showed noticeable signs of astigmatism or drift were discarded.

### Analysis and structure determination

The movie frames were aligned with MOTIONCORR (Li et al., 2013) for whole-image motion correction. Contrast transfer function parameters for the micrographs were estimated using Gctf (Zhang, 2016). Particles were picked using RELION (Scheres, 2012). References for template-based particle picking (Scheres, 2015) were obtained from 2D class averages that were calculated from particles picked with EMAN2 (Tang et al., 2007) from a subset of the micrographs. 2D class averaging, 3D classification and refinements were done using RELION-1.4 (Scheres, 2012). Both movie processing (Bai et al., 2013) in RELION-1.4 and particle ‘‘polishing’’ was performed for all selected particles for 3D refinement. Resolutions reported here are based on the gold-standard FSC = 0.143 criterion (Scheres and Chen, 2012). All maps were further processed for the modulation transfer function of the detector, and sharpened (Rosenthal and Henderson, 2003). Local resolution was estimated using Relion and ResMap (Kucukelbir et al., 2014).

A dataset of about 2100 images was recorded from two independent data acquisition sessions. An initial reconstruction was made from all selected particles (394,672) after 2D class averaging using the yeast 40S crystal structure (PDB: 4V88) low-pass filtered to 40 Å as an initial model. Next, a 3D classification into 8 classes with fine angular sampling and local searches was performed to remove bad particles/empty 40S particles from the data. Two highly populated classes (70%; 276,269 particles) showed density for the TC. In a second round of 3D classification into 4 classes, only one of the classes (157,868 particles, 40 % of the total; Map 1) still had clear density for the TC and corresponds to a closed conformation of the 48S with eIF5-NTD in place of eIF1, and yielded a resolution of 3.0 Å.

The map yielded a high overall resolution but poor local resolutions for peripheral and flexible elements like eIF3, eIF2β and eIF2γ, so we decided to collect an additional dataset in the presence of 1% formaldehyde. For this dataset about 1500 images were recorded, and 312,041 particles were selected after two-dimensional classification. After obtaining an initial three-dimensional refined model, two consecutive rounds of 3D classification with fine angular sampling and local searches were performed. In the second round of 3D classification, only 2 classes containing TC were selected (113,838 particles, 36.5 % of the total) and refined to high resolution after movie processing (3.6 Å). The model obtained is identical to that obtained with the non-crosslinked data although with higher occupancies for eIF3 and eIF2.

Then the particles from both datasets (non-crosslinked and crosslinked) were combined and we applied a strategy based on the reported method of masked 3D classifications with subtraction of the residual signal (Bai et al., 2015), by creating three different masks around the densities attributed to the PCI domains of eIF3, the eIF3bgi sub-module and the TC. We used ‘focused’ 3D classifications using these masks to isolate four distinct and well-defined maps, as follows. A) Map A, showing higher occupancy and best local resolution for eIF3 PCI domains [53,870 particles, 3.5Å], B) Map B showing higher occupancy and best local resolution for eIF3 bgi sub-module [54,699 particles, 3.5Å], and C) By using the TC mask for focused 3D classifications we obtained the following two different maps showing a different TC conformation as a result of the rotation of subunits eIF2γ, eIF2β and domain D3 of eIF2α around the acceptor arm of the tRNA_i_. 1) Map C1 presents the TC in conformation 1 [74,772 particles, 3.5Å], 2) Map C2 presents the TC in conformation 2, and density for eIF2β is weaker [137,103 particles, 3.1Å].

To ensure that the addition of the crosslinker to the latter dataset was not producing undesirable artifacts, each of these four classes was divided again into two subsets using the particles belonging to the crosslinked/non-crosslinked datasets and were refined independently; in all cases, the maps were identical to each other but at lower resolution than the one with all particles combined (see Table 3).

### Model building and refinement

In all five maps the conformations of 40S, eIF1A, eIF5-NTD, tRNA, mRNA and domains D1 and D2 of eIF2α are identical. Thus, modeling of all these elements was done in the higher resolution map (3.0Å, non-crosslinked dataset only; Map 1), whereas maps resulting from local masked classifications were essentially used for model building of the various subunits/domains of eIF3 and of eIF2β and eIF2γ. In more detail, the atomic model of py48S in closed conformation (PDB: 3JAP) was placed into density by rigid-body fitting using Chimera (Pettersen et al., 2004). Then the body and head of the 40S were independently fitted. Wild type tRNA_i_ was taken from PDB: 3JAQ for initial rigid-body fitting into its corresponding density. eIF3b subunit was taken from PDB: 4NOX and together with eIF3i and eIF3a C-term helix, placed into density by rigid-body fitting at the solvent side of the 40S in a position similar to that found previously (des Georges et al., 2015). Possible residue numbering for eIF3a C-term helix is based on eIF3a secondary structure predictions and its known interactions with eIF3b and the eIF3b RRM domain (Chiu et al., 2010). Next, the NMR structure of the N-terminal domain of eIF5 (PDB: 2E9H) from *Homo sapiens* was docked into density using Chimera. Then, each chain of the model (including ribosomal proteins, rRNA segments, protein factors and tRNA and mRNA) was rigid-body fitted in Coot (Emsley et al., 2010) and further model building was also done in Coot v0.8.

Model refinement was carried out in Refmac v5.8 optimized for electron microscopy (Brown et al., 2015), using external restraints generated by ProSMART and LIBG (Brown et al., 2015). All maps, including the one of highest resolution (Map 1) and maps A, B, C1 and C2 were used for refining. Average FSC was monitored during refinement. The final model was validated using MolProbity (Chen et al., 2010). Cross-validation against overfitting was calculated as previously described (Brown et al., 2015; Amunts et al., 2014). Refinement statistics for the last refinements, done in Map 1, are given in Table 1. All figures were generated using PyMOL (DeLano, 2006) Coot or Chimera.

### Plasmid constructions

Plasmids used in the present study are listed in Table S2. Plasmid pAS5-101 harboring *TIF5-FL* (Saini et al., 2014), encoding eIF5 tagged with FLAG epitope at the C-terminus, was used as template for constructing *TIF5* mutant alleles by fusion PCR. The resulting amplicons were inserted between the EcoRI and SalI sites of single-copy plasmid YCplac111, and the subcloned fragments of all mutant constructs were confirmed by DNA sequencing.

### Yeast strain construction

Yeast strains used in this study are listed in Table S2 and were constructed by introducing the plasmid-borne *TIF5* mutant alleles into the *tif5Δ his4-301* strain ASY100 containing the *TIF5, URA3* plasmid p3342, which was subsequently evicted by counter-selection on medium containing 5-fluoro-orotic acid.

### Biochemical assays with yeast extracts

Assays of β-galactosidase activity in whole cell extracts (WCEs) were performed as described previously (Moehle and Hinnebusch, 1991). For Western analysis, WCEs were prepared by trichloroacetic acid extraction as described previously (Reid and Schatz, 1982) and immunoblot analysis was conducted as described (Olsen et al., 2003) using antibodies against FLAG epitope (Sigma #F-3165) to detect eIF5-FL proteins, or against eIF2Bϵ/Gcd6 as a loading control (Bushman et al., 1993).

Assays of rates of eIF1A dissociation and P_i_ release from reconstituted 43S·mRNA PICs

The kinetics of eIF1A dissociation and P_i_ release from reconstituted PICs were conducted exactly as described in (Saini et al., 2014).

## Data Resources

Five maps have been deposited in the EMDB with accession codes EMDB: 4328, EMDB: 4330, EMDB: 4331, EMDB: 4327, EMDB: 4329, for the sample 1 map, Map A, Map B, Map C1 and Map C2, respectively. Two atomic coordinate models have been deposited in the PDB with accession codes PDB: 6FYX, PDB: 6FYY, for models showing TC in conformation 1 and conformation 2, respectively.

## Supplemental information

Supplemental information includes two tables and eight figures.

## Author Contributions

JLL and TH made the samples, collected and analyzed the data, determined the cryo-EM structures and wrote a first draft of the manuscript. YG prepared some of the reagents for the formation of cryo-EM grids. SK, RK and AKS performed the genetic experiments and helped write the manuscript. JN performed the biochemical experiments, and helped write the manuscript. JRL, AGH and VR supervised the work and helped to write the manuscript.

## ACKNOWLEDGEMENTS

We are grateful to CG Savva, G McMullan for technical support with cryo-EM, T Darling and J Grimmett for help with computing and J. Brasa for help with figures/movies. JL was supported by a FEBS postdoctoral fellowship.

This work was funded by grants to from the UK Medical Research Council (MC_U105184332), Wellcome Trust Senior Investigator award (WT096570), the Agouron Institute and the Jeantet Foundation to VR; by the Department of Science and Technology, Government of India Grant [Int/NZ/P-2/13] to AKS; from the NIH (GM62128) formerly to JRL; the Human Frontiers in Science Program (RGP-0028/2009) to AGH, JRL and VR; and by the Intramural Research Program of the NIH (AGH, JRL).

## Supplementary Figures and movies

**Figure 1 – figure supplement 1.**
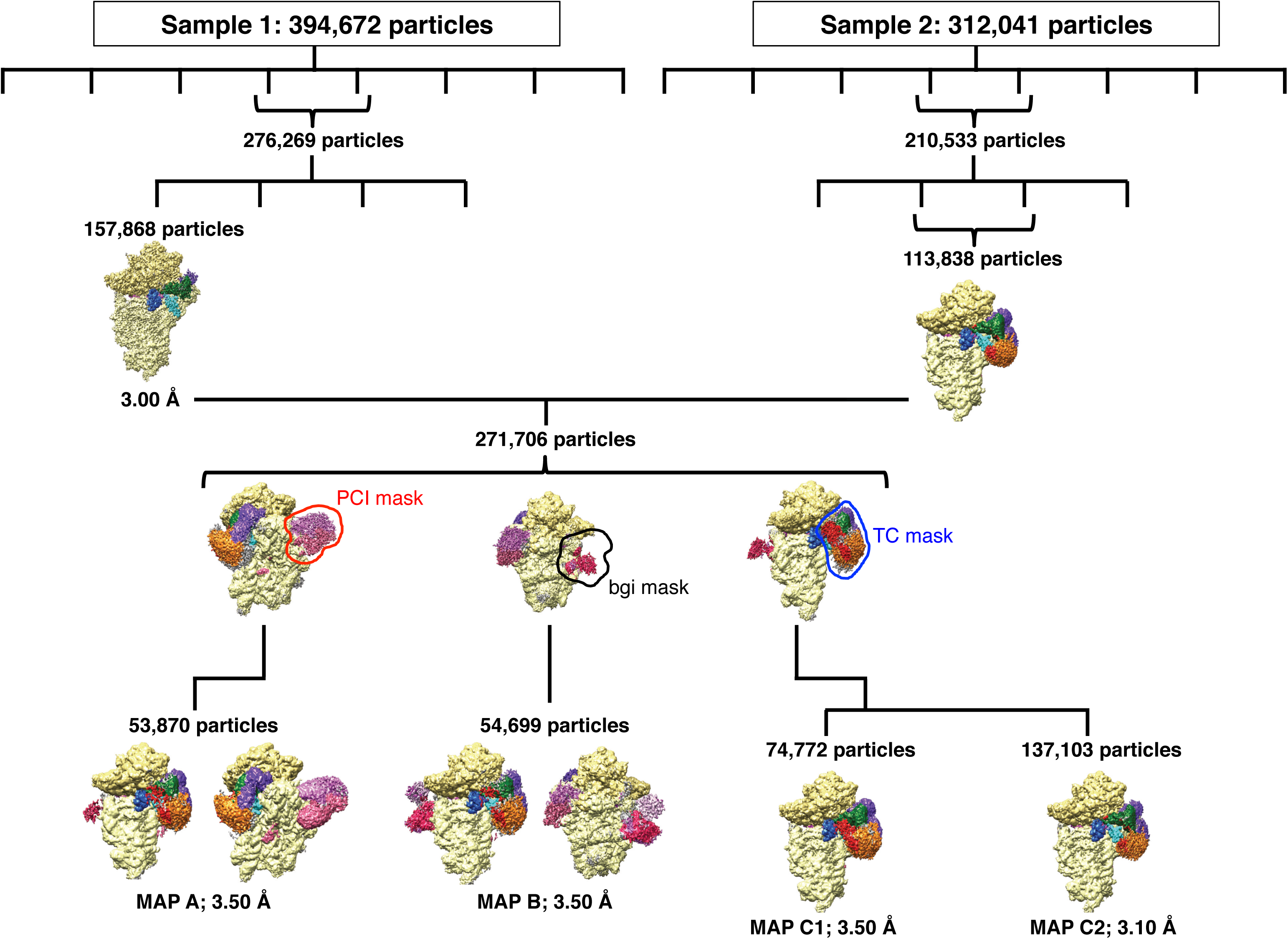
Scheme of 3D classification of data. For Sample 1 (un-crosslinked dataset) 394,672 particles were selected after 2D classification and an initial 3D refinement was done. After two rounds of 3D-classification, a class containing clear density for TC was refined to 3.0 Å-resolution (Map 1). Densities for peripheral and flexible elements like eIF3, eIF2β and eIF2γ are not seen in the completely sharpened map. For Sample 2 (1%-formaldehyde-crosslinked dataset) 312,041 particles were selected after 2D classification and an initial 3D refinement was done. After two rounds of 3D-classification, a class containing clear density for TC was obtained. Particles from both samples were joined and further separate, focused 3D classifications were carried out. The eIF3 masks ‘PCI mask’ and ‘bgi mask’ as well as the ‘TC mask’ used separately for focused 3D classification on the combined 271,706 particles are shown by an outline. The scheme shows how each map (Maps: 1, A, B, C1 and C2) was obtained. See ‘Analysis and structure determination’ section for detailed information.

**Figure 1 – figure supplement 2.**
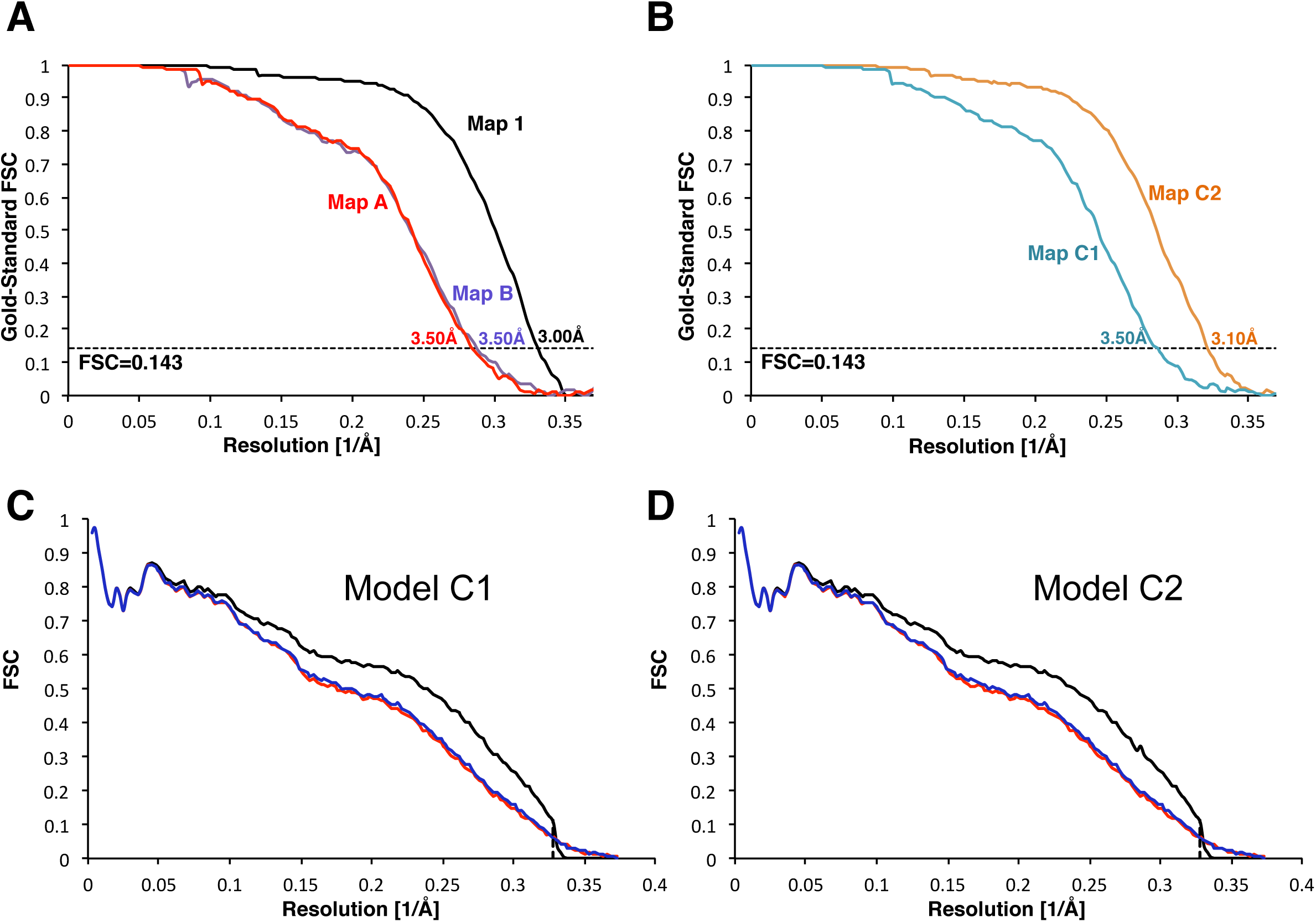
Validation of the maps. (A) Gold-standard Fourier Shell Correlation (FSC) curves for the Maps 1, A and B. (B) Gold-standard Fourier Shell Correlation (FSC) curves for Maps C1 and C2. (C) Analysis of overfitting by cross-validation of the Model C1. FSC_work_ curves (red) corresponding to the refined model versus the half-map it was refined against, and FSC_test_ curves (green), i.e. those calculated between the refined atomic model and the other half-map. The black curve shows the FSC curve between a reconstruction from all particles and the model refined against the map. (D) Analysis of overfitting by cross-validation, similar to that in (C), of the Model C2.

**Figure 1 – figure supplement 3.**
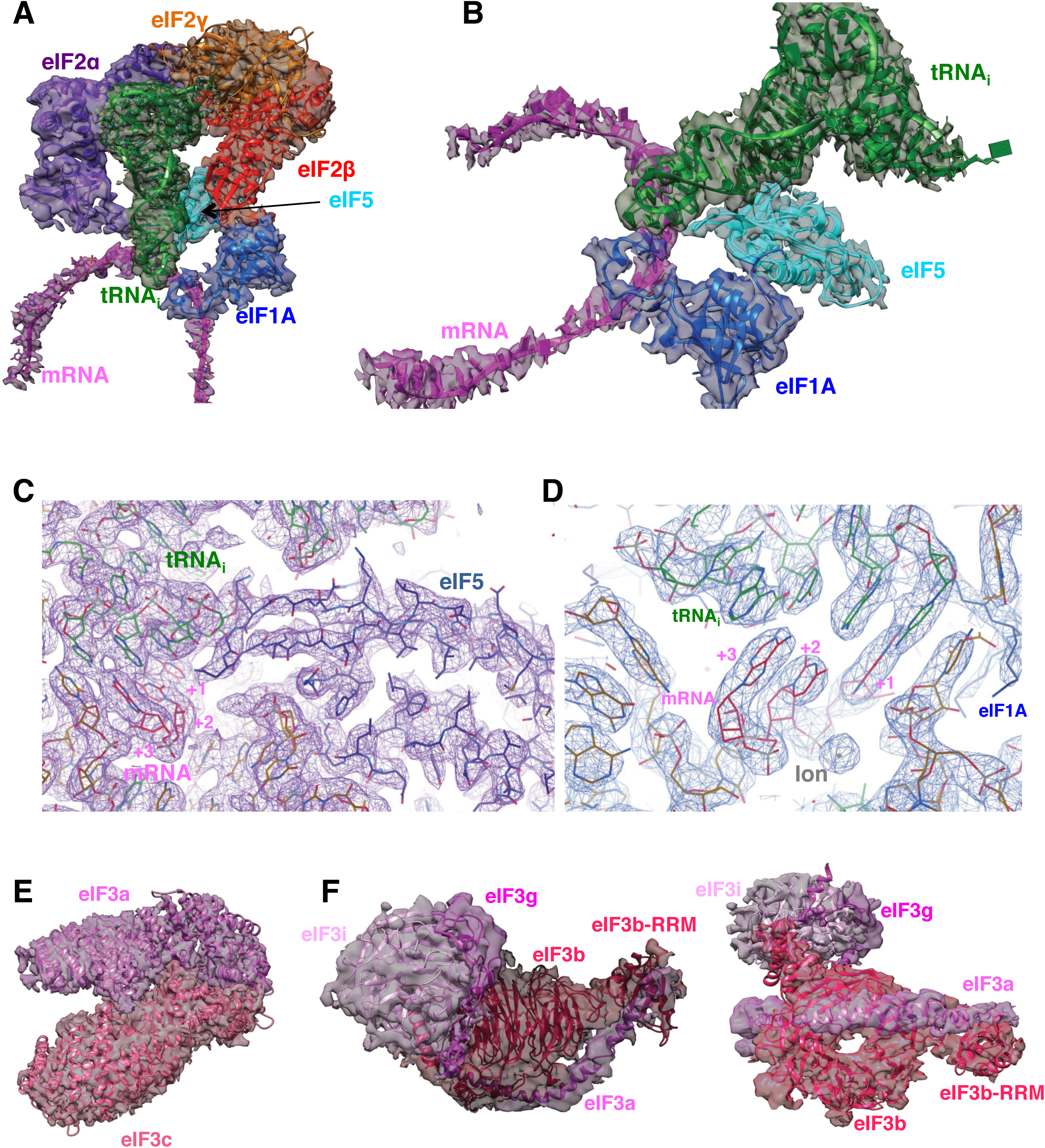
Fitting of ligands in density maps. (A) Fitting of mRNA (magenta), eIF1A (marine), eIF5-NTD (cyan), tRNA_i_ (green), eIF2α (purple), eIF2β (red) and eIF2γ (orange) in Map C1. (B) As in A, but shown in a different orientation and omitting eIF2 components to highlight the fitting of eIF5-NTD. (C) High-resolution features for tRNA_i_, mRNA and eIF5-NTD at the P site, at 3.0 Å-resolution (Map 1) (D) As in C, but showing the ion centered around the mRNA codon (E) Fitting of the eIF3a/c PCI domain dimer in map A. (F) Fitting of the eIF3bgi subcomplex in map B, shown in two orientations.

**Figure 1 – figure supplement 4.**
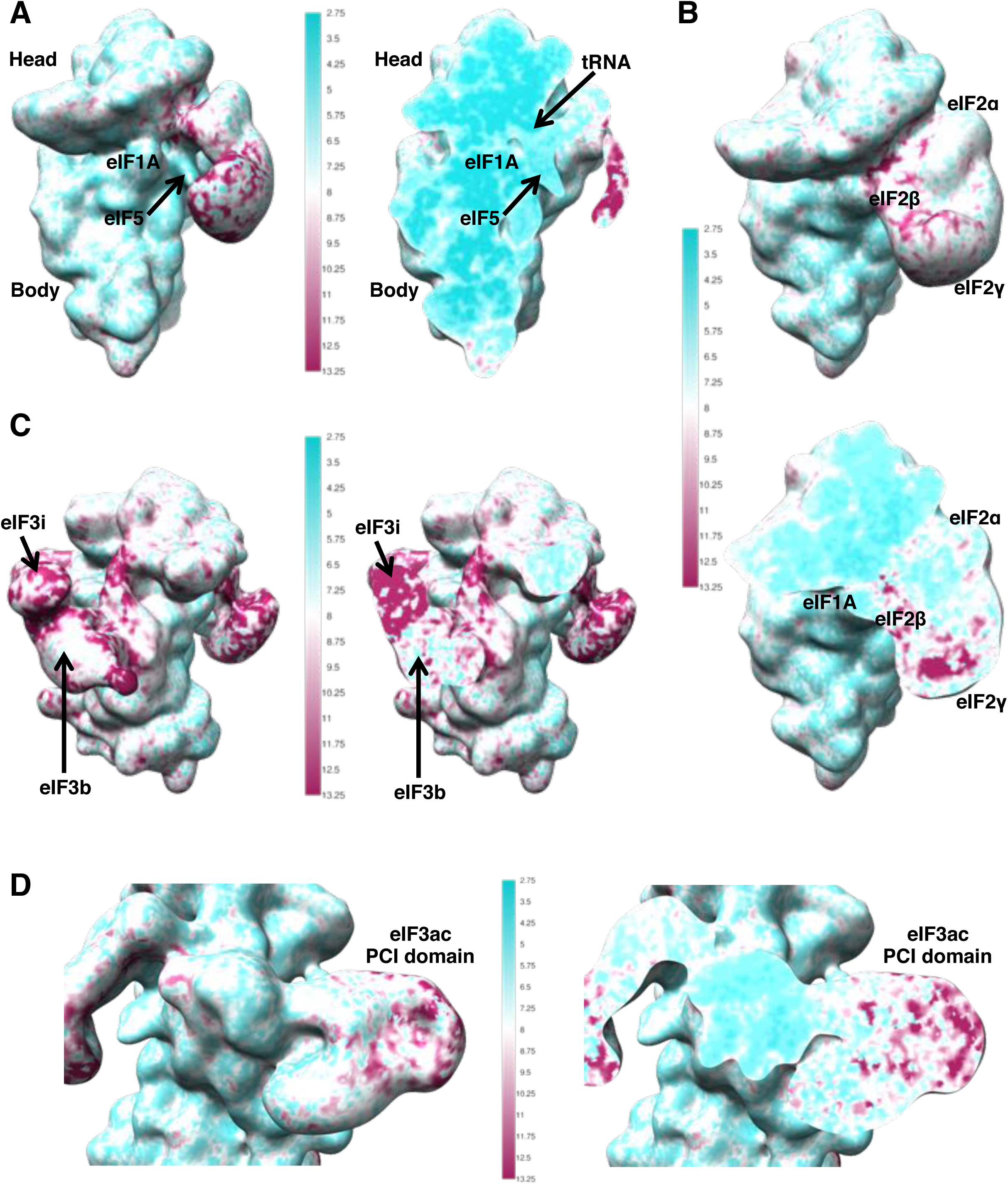
Map quality and local resolution. Surface (left or top) and cross-sections (right or bottom) of gaussian-filtered maps, colored according to local resolution. (A) Map 1 (B) Map C1 (C) Map B (D) Map A

**Figure 1 – figure supplement 5.**
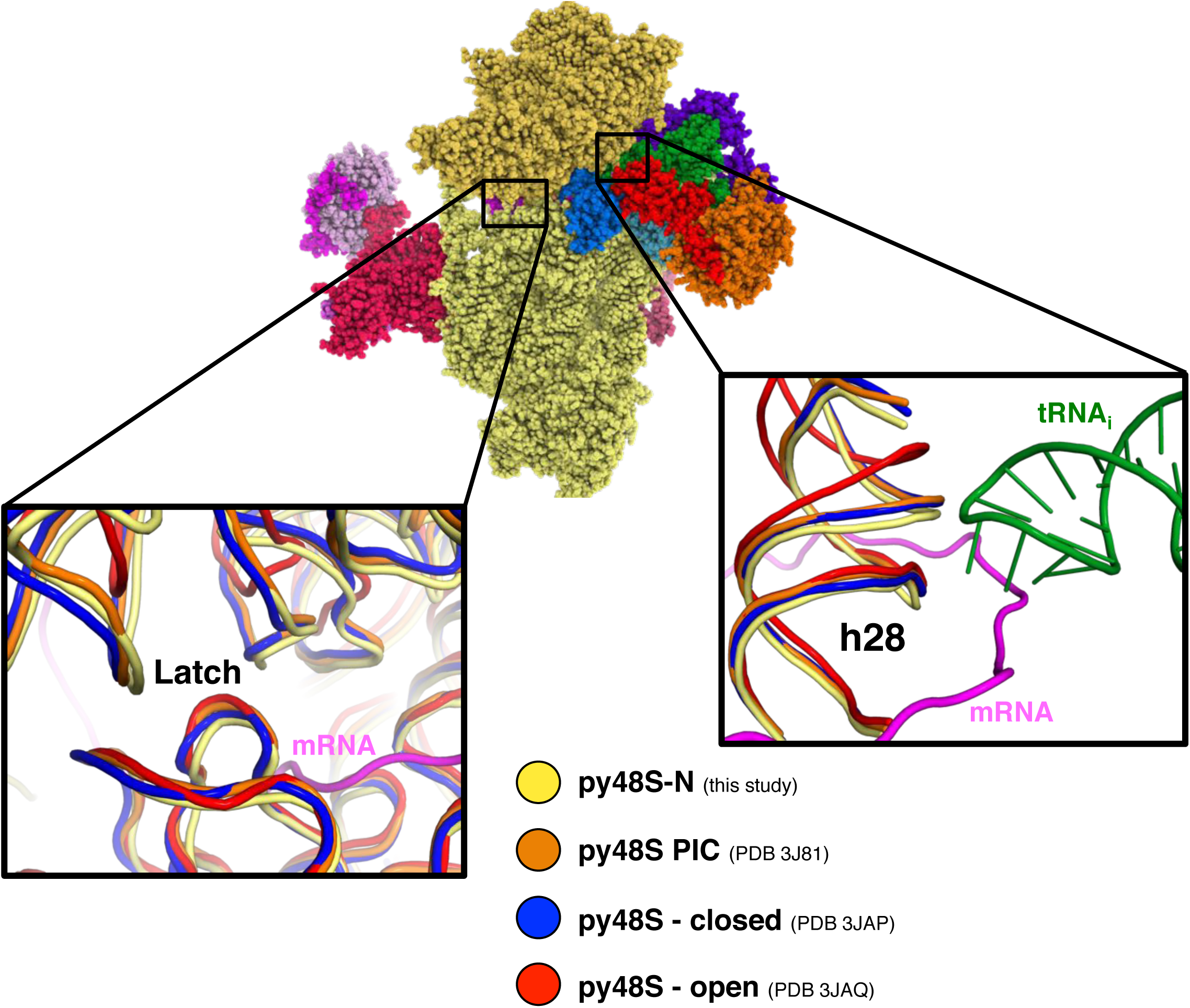
Latch and h28 conformation and head closure in different py48S PICs. Superimposition of different py48S complexes shows a different degree of head closure around the latch area (left inset). Right inset shows distinct h28 conformations; only py48S-open presents a non-compressed h28 whereas in all other PICs, including py48S-eIF5N presented here, h28 is compressed.

**Figure 2 – figure supplement 1.**
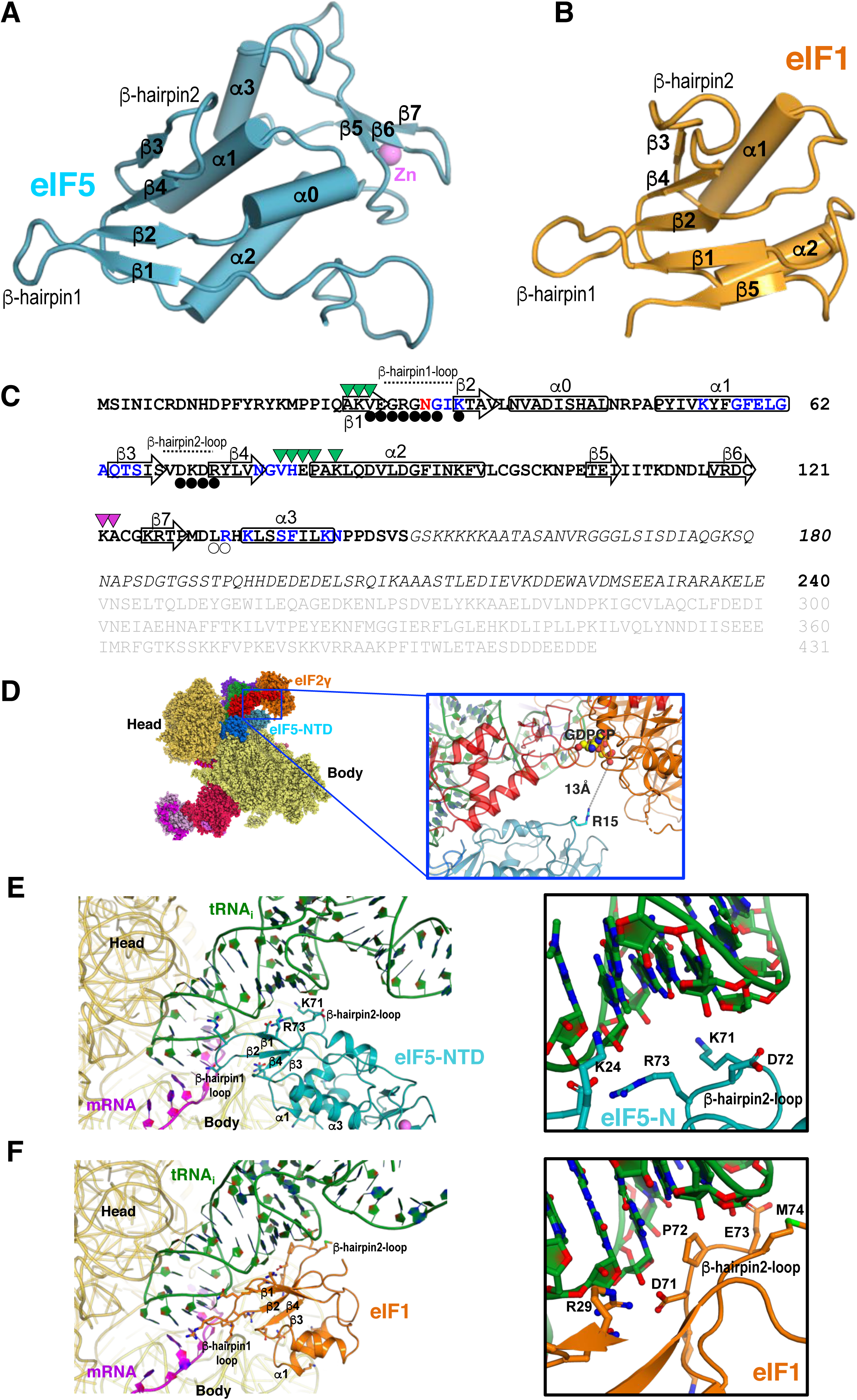
eIF5 and eIF1 comparison. (A) Cartoon representation of eIF5 (this study). Secondary structure elements are labeled. (B) Cartoon representation of eIF1 (from PDB:3J81) with labeled secondary structure elements. (C) Amino acid sequence and topology of secondary structure elements of eIF5. The β-strands and α-helices are depicted as superimposed arrows and rectangles, respectively. Residues belonging to the eIF5-NTD are in bold whereas those in light grey (comprising the eIF5-CTD) or in italics (comprising the linker connecting the NTD and CTD) are not visible in the present structure. In eIF5-NTD, the residues having decreased accessibility upon binding to 40S (in blue), tRNA (black circles), eIF2β (transparent circles), eIF2γ (violet triangles) and eIF1A (green triangles) in py48S-eIF5N are indicated. Asn30 interacting with mRNA, tRNA and the 40S simultaneously is shown in red. (D) Cartoon representation showing how Arg15 (essential for GAP activity) is 13Å away from bound nucleotide in eIF2γ. (E) Contacts of eIF5-NTD with tRNA_i_ or mRNA in py48S-eIF5N. Only secondary structure elements involved in the contacts are labeled. Two basic residues located in β-hairpin loop-2 of eIF5-NTD (K71, R73) interact with tRNA_i_. (F) Contacts of eIF1 with tRNA_i_ or mRNA in the py48S complex (from PDB:3J81). Only secondary structure elements involved in the contacts are labeled, as in (E). The analogous secondary structure elements participate in the interactions in the two proteins; however, eIF1 β-hairpin loop-2 is negatively charged and exerts electrostatic repulsion between eIF1 and the tRNA_i_ rather than the attraction afforded by basic residues in the eIF5-NTD loop-2 (E).

**Figure 2 – figure supplement 2.**
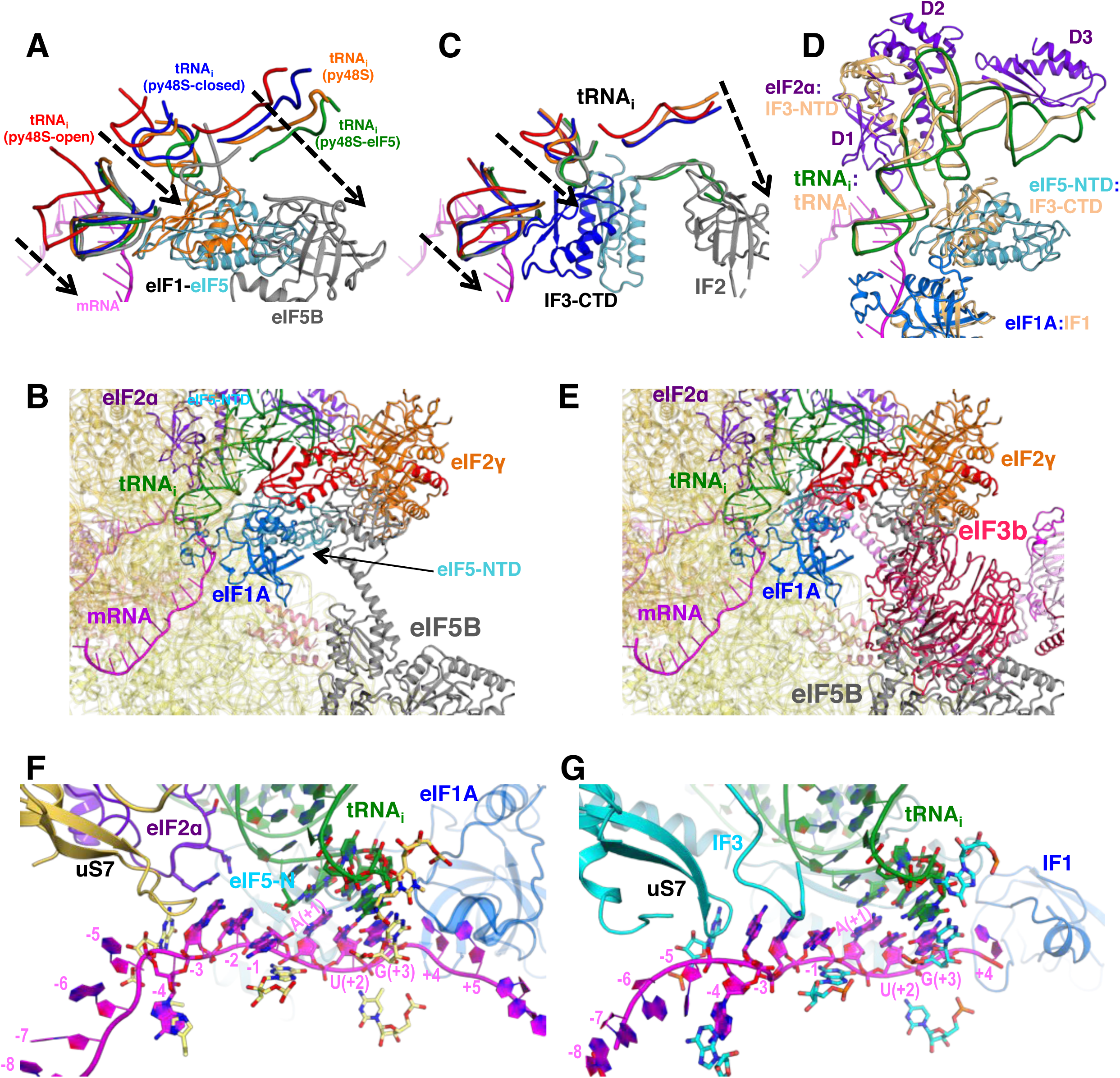
Comparison of eukaryotic and bacterial initiation following start codon recognition. (A) Accommodation of tRNA_i_ in the P site in different yeast PICs. Similar representation to that in Figure 2C but including eIF5B and its associated tRNA (in gray) from the structure of an 80S initiation complex (PDB:4UJD) instead of bacterial IF2/tRNA_i_ from bacterial PIC-III. For simplicity, only nucleotides 7-15 (D-loop) +26-43 (ASL) +69-76 (acceptor arm) from tRNA are shown. (B) Same orientation as in (A) of the superimposition of different bacterial PICs. When comparing panels C and D, we observe a similar tRNA accommodation process which depends on the displacement of eIF1/IF3 from near the P site. The present structure with eIF5-NTD in place of eIF1 could be considered the eukaryotic counterpart of bacterial PIC-4 (tRNA in green; IF3-CTD in cyan) or PIC-III (tRNA in gray; IF3-CTD in cyan) wherein IF3-CTD is completely displaced from the P site, since in both prokaryotic/eukaryotic PIC structures the tRNA accommodation process is very similar. (C) Similarity between locations and interactions of different eukaryotic and bacterial initiation factors and tRNA_i_s in closed PIC structures. Bacterial PIC-3 (in yellow) and the py48S-eIF5N structures were superimposed by aligning the 40S/30S bodies. The tRNA_i_s superpose quite well, and IF1:eIF1A, IF3-CTD:eIF5-NTD (or eIF1, not shown), IF3-NTD:eIF2α (domains D1 and D2) all occupy similar positions in two complexes. (D) Superimposition of eIF5B from the 80S initiation complex (PDB:4UJD) (in gray) with py48S-eIF5N shows a potential coexistence of eIF5B and eIF5-NTD in the same PIC, as eIF5B would clash with eIF2 but not with eIF5-NTD. (E) Superimposition of eIF5B from the 80S initiation complex (PDB:4UJD) with py48S-closed (PDB 6GSN) shows the incompatibility of eIF3 bound at the subunit interface with eIF5B, based on predicted extensive clashing of eIF3 subunits with eIF5B. (F) Similar representation to that in Figure 6C of the mRNA at the P site and flanking regions of py48S-eIF5N. (G) Same orientation as in (F) of the equivalent region of a bacterial PIC in closed conformation (PIC4; 5LMU), revealing an almost identical mRNA configuration and equivalent elements stabilizing the mRNA as observed in (F).

**Figure 3 – figure supplement 1.**
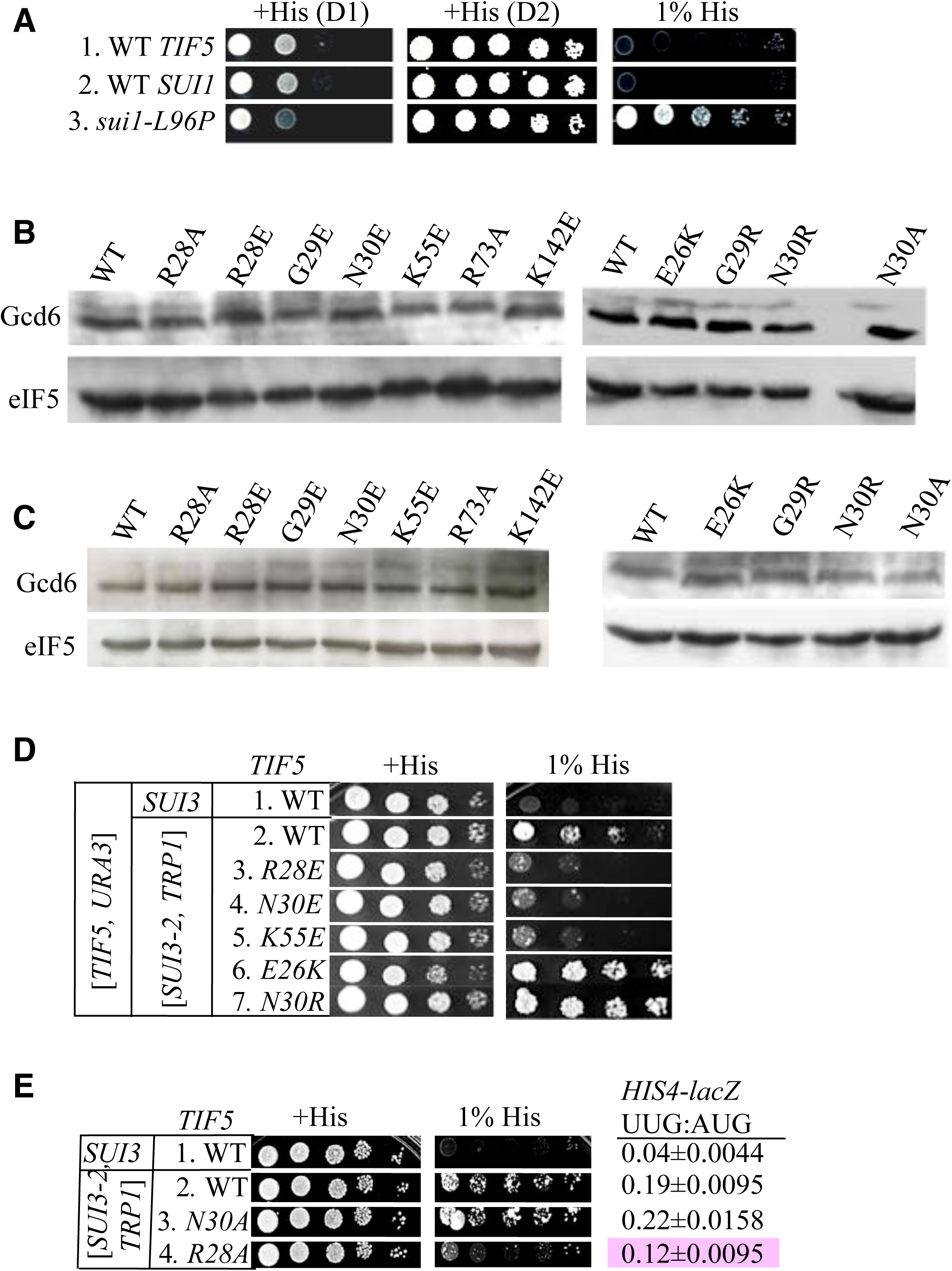
(A) Strains ASY101, PMY30 and PMY33 with the indicated relevant genotypes (rows 1-3, respectively) were spotted on SC-L medium supplemented with 0.3 mM histidine (+His) or 0.0003 mM histidine (1% His), and incubated for 1d or 2d (+His) or 6d (1% His) at 30°C. (B-C) Ssu^−^ and Sui^−^ substitutions do not alter expression of FLAG-tagged eIF5 proteins. Derivatives of *his4-301 tif5Δ* strain ASY100 containing the indicated *TIF5-FL* alleles on sc *LEU2* plasmids and harboring only chromosomal WT *SUI3* (B) or also plasmid-borne *SUI3-2* (C) were cultured in SC medium lacking leucine (B) or leucine and tryptophan (C). WCEs were prepared using trichloroacetic acid, resolved by SDS-PAGE and immunoblotted using antibodies against FLAG epitope or Gcd6. (D) Dominant Slg^−^ and His^+^/Sui^−^ phenotypes were determined for derivatives of *his4*-*301 tif5Δ* strain ASY100 harboring WT *TIF5, URA3* plasmid p3342, the indicated *TIF5-FL* alleles on *LEU2* plasmids, and either *SUI3-2* plasmid pRSSUI3-S264Y-W (rows 2–7) or empty vector (row 1). Ten-fold serial dilutions were spotted on SC media lacking leucine, tryptophan and uracil supplemented either with 0.3 mM histidine (+His) or 0.0003 mM histidine (1% His) and incubated for 3d (+His) or 6d (1% His) at 30°C. (E) Strains harbouring the indicated *TIF5-FL* alleles and either *SUI3-2* plasmid pRSSUI3-S264Y-W (rows 2-4) or empty vector (row 1) were analyzed as in (D) except that cells were spotted on SC media lacking leucine and tryptophan. *HIS4-lacZ* initiation ratios were determined in these strains as in Figure 3C. The Ssu^−^ phenotype of *-R28A* is highlighted in pink.

**Movie 1. Movie highlighting tRNA accommodation, leading to eIF1 dissociation and eIF5-NTD recruitment to the 48S complex.** Detailed contacts of eIF5-NTD with other elements of the 48S complex are also shown.

## References

Acker, M. G., Kolitz, S. E., Mitchell, S. F., Nanda, J. S., and Lorsch, J. R. (2007). Reconstitution of yeast translation initiation. Methods Enzymol 430, 111–145.

Aitken, C. E., Beznosková, P., Vlčkova, V., Chiu, W. L., Zhou, F., Valášek, L. S., Hinnebusch, A. G., and Lorsch, J. R. (2016). Eukaryotic translation initiation factor 3 plays distinct roles at the mRNA entry and exit channels of the ribosomal preinitiation complex. Elife 5,

Algire, M. A., Maag, D., and Lorsch, J. R. (2005). Pi release from eIF2, not GTP hydrolysis, is the step controlled by start-site selection during eukaryotic translation initiation. Mol Cell 20, 251–262.

Amunts, A., Brown, A., Bai, X. C., Llacer, J. L., Hussain, T., Emsley, P., Long, F., Murshudov, G., Scheres, S. H., and Ramakrishnan, V. (2014). Structure of the yeast mitochondrial large ribosomal subunit. Science 343, 1485–1489.

Aylett, C. H., and Ban, N. (2017). Eukaryotic aspects of translation initiation brought into focus. Philos Trans R Soc Lond B Biol Sci 372,

Aylett, C. H., Boehringer, D., Erzberger, J. P., Schaefer, T., and Ban, N. (2015). Structure of a Yeast 40S-eIF1-eIF1A-eIF3-eIF3j initiation complex. Nat Struct Mol Biol

Bai, X. C., Fernandez, I. S., McMullan, G., and Scheres, S. H. (2013). Ribosome structures to near-atomic resolution from thirty thousand cryo-EM particles. Elife 2, e00461.

Bai, X. C., Rajendra, E., Yang, G., Shi, Y., and Scheres, S. H. (2015). Sampling the conformational space of the catalytic subunit of human γ-secretase. Elife 4,

Brown, A., Long, F., Nicholls, R. A., Toots, J., Emsley, P., and Murshudov, G. (2015). Tools for macromolecular model building and refinement into electron cryo-microscopy reconstructions. Acta Crystallogr D Biol Crystallogr 71, 136–153.

Bushman, J. L., Foiani, M., Cigan, A. M., Paddon, C. J., and Hinnebusch, A. G. (1993). Guanine nucleotide exchange factor for eukaryotic translation initiation factor 2 in Saccharomyces cerevisiae: interactions between the essential subunits GCD2, GCD6, and GCD7 and the regulatory subunit GCN3. Mol Cell Biol 13, 4618–4631.

Chen, V. B., Arendall, W. B., Headd, J. J., Keedy, D. A., Immormino, R. M., Kapral, G. J., Murray, L. W., Richardson, J. S., and Richardson, D. C. (2010). MolProbity: all-atom structure validation for macromolecular crystallography. Acta Crystallogr D Biol Crystallogr 66, 12–21.

Cheung, Y. N., Maag, D., Mitchell, S. F., Fekete, C. A., Algire, M. A., Takacs, J. E., Shirokikh, N., Pestova, T., Lorsch, J. R., and Hinnebusch, A. G. (2007). Dissociation of eIF1 from the 40S ribosomal subunit is a key step in start codon selection in vivo. Genes Dev 21, 1217–1230.

Chiu, W. L., Wagner, S., Herrmannova, A., Burela, L., Zhang, F., Saini, A. K., Valasek, L., and Hinnebusch, A. G. (2010). The C-terminal region of eukaryotic translation initiation factor 3a (eIF3a) promotes mRNA recruitment, scanning, and, together with eIF3j and the eIF3b RNA recognition motif, selection of AUG start codons. Mol Cell Biol 30, 4415–4434.

Conte, M. R., Kelly, G., Babon, J., Sanfelice, D., Youell, J., Smerdon, S. J., and Proud, C. G. (2006). Structure of the eukaryotic initiation factor (eIF) 5 reveals a fold common to several translation factors. Biochemistry 45, 4550–4558.

Cuchalova, L., Kouba, T., Herrmannova, A., Danyi, I., Chiu, W. L., and Valasek, L. (2010). The RNA recognition motif of eukaryotic translation initiation factor 3g (eIF3g) is required for resumption of scanning of posttermination ribosomes for reinitiation on GCN4 and together with eIF3i stimulates linear scanning. Mol Cell Biol 30, 4671–4686.

Das, S., Ghosh, R., and Maitra, U. (2001). Eukaryotic translation initiation factor 5 functions as a GTPase-activating protein. J Biol Chem 276, 6720–6726.

DeLano, W. L. (2006). The PyMOL Molecular Graphics System. http://www.pymol.org

des Georges, A., Dhote, V., Kuhn, L., Hellen, C. U., Pestova, T. V., Frank, J., and Hashem, Y. (2015). Structure of mammalian eIF3 in the context of the 43S preinitiation complex. Nature 525, 491–495.

Dong, J., Aitken, C. E., Thakur, A., Shin, B. S., Lorsch, J. R., and Hinnebusch, A. G. (2017). Rps3/uS3 promotes mRNA binding at the 40S ribosome entry channel and stabilizes preinitiation complexes at start codons. Proc Natl Acad Sci U S A 114, E2126–E2135.

Emsley, P., Lohkamp, B., Scott, W. G., and Cowtan, K. (2010). Features and development of Coot. Acta Crystallogr D Biol Crystallogr 66, 486–501.

Fekete, C. A., Mitchell, S. F., Cherkasova, V. A., Applefield, D., Algire, M. A., Maag, D., Saini, A. K., Lorsch, J. R., and Hinnebusch, A. G. (2007). N-and C-terminal residues of eIF1A have opposing effects on the fidelity of start codon selection. EMBO J 26, 1602–1614.

Fernandez, I. S., Bai, X. C., Hussain, T., Kelley, A. C., Lorsch, J. R., Ramakrishnan, V., and Scheres, S. H. (2013). Molecular architecture of a eukaryotic translational initiation complex. Science 342, 1240585.

Fernandez, I. S., Bai, X. C., Murshudov, G., Scheres, S. H., and Ramakrishnan, V. (2014). Initiation of translation by cricket paralysis virus IRES requires its translocation in the ribosome. Cell 157, 823–831.

Hashem, Y., des Georges, A., Dhote, V., Langlois, R., Liao, H. Y., Grassucci, R. A., Hellen, C. U., Pestova, T. V., and Frank, J. (2013). Structure of the mammalian ribosomal 43S preinitiation complex bound to the scanning factor DHX29. Cell 153, 1108–1119.

Hinnebusch, A. G. (2014). The scanning mechanism of eukaryotic translation initiation. Annu Rev Biochem 83, 779–812.

Hinnebusch, A. G. (2017). Structural Insights into the Mechanism of Scanning and Start Codon Recognition in Eukaryotic Translation Initiation. Trends Biochem Sci 42, 589–611.

Huang, H. K., Yoon, H., Hannig, E. M., and Donahue, T. F. (1997). GTP hydrolysis controls stringent selection of the AUG start codon during translation initiation in Saccharomyces cerevisiae. Genes Dev 11, 2396–2413.

Hussain, T., Llacer, J. L., Fernandez, I. S., Munoz, A., Martin-Marcos, P., Savva, C. G., Lorsch, J. R., Hinnebusch, A. G., and Ramakrishnan, V. (2014). Structural changes enable start codon recognition by the eukaryotic translation initiation complex. Cell 159, 597–607.

Hussain, T., Llácer, J. L., Wimberly, B. T., Kieft, J. S., and Ramakrishnan, V. (2016). Large-Scale Movements of IF3 and tRNA during Bacterial Translation Initiation. Cell 167, 133–144.e13.

Ivanov, I. P., Loughran, G., Sachs, M. S., and Atkins, J. F. (2010). Initiation context modulates autoregulation of eukaryotic translation initiation factor 1 (eIF1). Proc Natl Acad Sci U S A 107, 18056–18060.

Kapp, L. D., and Lorsch, J. R. (2004). GTP-dependent recognition of the methionine moiety on initiator tRNA by translation factor eIF2. J Mol Biol 335, 923–936.

Kirillov, S. V., Wower, J., Hixson, S. S., and Zimmermann, R. A. (2002). Transit of tRNA through the Escherichia coli ribosome: cross-linking of the 3’ end of tRNA to ribosomal proteins at the P and E sites. FEBS Lett 514, 60–66.

Kozak, M. (1986). Point mutations define a sequence flanking the AUG initiator codon that modulates translation by eukaryotic ribosomes. Cell 44, 283–292.

Kozel, C., Thompson, B., Hustak, S., Moore, C., Nakashima, A., Singh, C. R., Reid, M., Cox, C., Papadopoulos, E., Luna, R. E., Anderson, A., Tagami, H., Hiraishi, H., Slone, E. A., Yoshino, K. I., Asano, M., Gillaspie, S., Nietfeld, J., Perchellet, J. P., Rothenburg, S., Masai, H., Wagner, G., Beeser, A., Kikkawa, U., Fleming, S. D., and Asano, K. (2016). Overexpression of eIF5 or its protein mimic 5MP perturbs eIF2 function and induces ATF4 translation through delayed re-initiation. Nucleic Acids Res 44, 8704–8713.

Kucukelbir, A., Sigworth, F. J., and Tagare, H. D. (2014). Quantifying the local resolution of cryo-EM density maps. Nat Methods 11, 63–65.

Li, X., Mooney, P., Zheng, S., Booth, C. R., Braunfeld, M. B., Gubbens, S., Agard, D. A., and Cheng, Y. (2013). Electron counting and beam-induced motion correction enable near-atomic-resolution single-particle cryo-EM. Nat Methods 10, 584–590.

Llácer, J. L., Hussain, T., Marler, L., Aitken, C. E., Thakur, A., Lorsch, J. R., Hinnebusch, A. G., and Ramakrishnan, V. (2015). Conformational Differences between Open and Closed States of the Eukaryotic Translation Initiation Complex. Mol Cell 59, 399–412.

Lomakin, I. B., and Steitz, T. A. (2013). The initiation of mammalian protein synthesis and mRNA scanning mechanism. Nature 500, 307–311.

Loughran, G., Sachs, M. S., Atkins, J. F., and Ivanov, I. P. (2012). Stringency of start codon selection modulates autoregulation of translation initiation factor eIF5. Nucleic Acids Res 40, 2898–2906.

Maag, D., Algire, M. A., and Lorsch, J. R. (2006). Communication between eukaryotic translation initiation factors 5 and 1A within the ribosomal pre-initiation complex plays a role in start site selection. J Mol Biol 356, 724–737.

Maag, D., Fekete, C. A., Gryczynski, Z., and Lorsch, J. R. (2005). A conformational change in the eukaryotic translation preinitiation complex and release of eIF1 signal recognition of the start codon. Mol Cell 17, 265–275.

Martin-Marcos, P., Cheung, Y. N., and Hinnebusch, A. G. (2011). Functional elements in initiation factors 1, 1A, and 2beta discriminate against poor AUG context and non-AUG start codons. Mol Cell Biol 31, 4814–4831.

Martin-Marcos, P., Nanda, J., Luna, R. E., Wagner, G., Lorsch, J. R., and Hinnebusch, A. G. (2013). beta-Hairpin loop of eukaryotic initiation factor 1 (eIF1) mediates 40 S ribosome binding to regulate initiator tRNA(Met) recruitment and accuracy of AUG selection in vivo. J Biol Chem 288, 27546–27562.

Martin-Marcos, P., Nanda, J. S., Luna, R. E., Zhang, F., Saini, A. K., Cherkasova, V. A., Wagner, G., Lorsch, J. R., and Hinnebusch, A. G. (2014). Enhanced eIF1 binding to the 40S ribosome impedes conformational rearrangements of the preinitiation complex and elevates initiation accuracy. RNA 20, 150–167.

Martin-Marcos, P., Zhou, F., Karunasiri, C., Zhang, F., Dong, J., Nanda, J., Kulkarni, S. D., Sen, N. D., Tamame, M., Zeschnigk, M., Lorsch, J. R., and Hinnebusch, A. G. (2017). eIF1A residues implicated in cancer stabilize translation preinitiation complexes and favor suboptimal initiation sites in yeast. Elife 6,

Meyer, B., Wurm, J. P., Kötter, P., Leisegang, M. S., Schilling, V., Buchhaupt, M., Held, M., Bahr, U., Karas, M., Heckel, A., Bohnsack, M. T., Wöhnert, J., and Entian, K. D. (2011). The Bowen-Conradi syndrome protein Nep1 (Emg1) has a dual role in eukaryotic ribosome biogenesis, as an essential assembly factor, and in the methylation of Ψ1191 in yeast 18S rRNA. Nucleic Acids Res 39, 1526–1537.

Mitchell, S. F., Walker, S. E., Algire, M. A., Park, E. H., Hinnebusch, A. G., and Lorsch, J. R. (2010). The 5’-7-methylguanosine cap on eukaryotic mRNAs serves both to stimulate canonical translation initiation and to block an alternative pathway. Mol Cell 39, 950–962.

Moehle, C. M., and Hinnebusch, A. G. (1991). Association of RAP1 binding sites with stringent control of ribosomal protein gene transcription in Saccharomyces cerevisiae. Mol Cell Biol 11, 2723–2735.

Nanda, J. S., Cheung, Y. N., Takacs, J. E., Martin-Marcos, P., Saini, A. K., Hinnebusch, A. G., and Lorsch, J. R. (2009). eIF1 controls multiple steps in start codon recognition during eukaryotic translation initiation. J Mol Biol 394, 268–285.

Nanda, J. S., Saini, A. K., Munoz, A. M., Hinnebusch, A. G., and Lorsch, J. R. (2013). Coordinated movements of eukaryotic translation initiation factors eIF1, eIF1A, and eIF5 trigger phosphate release from eIF2 in response to start codon recognition by the ribosomal preinitiation complex. J Biol Chem 288, 5316–5329.

Olsen, D. S., Savner, E. M., Mathew, A., Zhang, F., Krishnamoorthy, T., Phan, L., and Hinnebusch, A. G. (2003). Domains of eIF1A that mediate binding to eIF2, eIF3 and eIF5B and promote ternary complex recruitment in vivo. EMBO J 22, 193–204.

Passmore, L. A., Schmeing, T. M., Maag, D., Applefield, D. J., Acker, M. G., Algire, M. A., Lorsch, J. R., and Ramakrishnan, V. (2007). The eukaryotic translation initiation factors eIF1 and eIF1A induce an open conformation of the 40S ribosome. Mol Cell 26, 41–50.

Paulin, F. E., Campbell, L. E., O’Brien, K., Loughlin, J., and Proud, C. G. (2001). Eukaryotic translation initiation factor 5 (eIF5) acts as a classical GTPase-activator protein. Curr Biol 11, 55–59.

Pettersen, E. F., Goddard, T. D., Huang, C. C., Couch, G. S., Greenblatt, D. M., Meng, E. C., and Ferrin, T. E. (2004). UCSF Chimera-a visualization system for exploratory research and analysis. J Comput Chem 25, 1605–1612.

Pisarev, A. V., Kolupaeva, V. G., Pisareva, V. P., Merrick, W. C., Hellen, C. U., and Pestova, T. V. (2006). Specific functional interactions of nucleotides at key -3 and +4 positions flanking the initiation codon with components of the mammalian 48S translation initiation complex. Genes Dev 20, 624–636.

Pisarev, A. V., Kolupaeva, V. G., Yusupov, M. M., Hellen, C. U., and Pestova, T. V. (2008). Ribosomal position and contacts of mRNA in eukaryotic translation initiation complexes. EMBO J 27, 1609–1621.

Pisareva, V. P., and Pisarev, A. V. (2014). eIF5 and eIF5B together stimulate 48S initiation complex formation during ribosomal scanning. Nucleic Acids Res 42, 12052–12069.

Rabl, J., Leibundgut, M., Ataide, S. F., Haag, A., and Ban, N. (2011). Crystal Structure of the Eukaryotic 40S Ribosomal Subunit in Complex with Initiation Factor 1. Science 331, 730–736.

Reid, G. A., and Schatz, G. (1982). Import of proteins into mitochondria. Yeast cells grown in the presence of carbonyl cyanide m-chlorophenylhydrazone accumulate massive amounts of some mitochondrial precursor polypeptides. J Biol Chem 257, 13056–13061.

Rosenthal, P. B., and Henderson, R. (2003). Optimal determination of particle orientation, absolute hand, and contrast loss in single-particle electron cryomicroscopy. J Mol Biol 333, 721–745.

Saini, A. K., Nanda, J. S., Martin-Marcos, P., Dong, J., Zhang, F., Bhardwaj, M., Lorsch, J. R., and Hinnebusch, A. G. (2014). Eukaryotic translation initiation factor eIF5 promotes the accuracy of start codon recognition by regulating Pi release and conformational transitions of the preinitiation complex. Nucleic Acids Res 42, 9623–9640.

Scheres, S. H. (2012). RELION: implementation of a Bayesian approach to cryo-EM structure determination. J Struct Biol 180, 519–530.

Scheres, S. H. (2015). Semi-automated selection of cryo-EM particles in RELION-1.3. J Struct Biol 189, 114–122.

Scheres, S. H., and Chen, S. (2012). Prevention of overfitting in cryo-EM structure determination. Nat Methods 9, 853–854.

Simonetti, A., Brito Querido, J., Myasnikov, A. G., Mancera-Martinez, E., Renaud, A., Kuhn, L., and Hashem, Y. (2016). eIF3 Peripheral Subunits Rearrangement after mRNA Binding and Start-Codon Recognition. Mol Cell 63, 206–217.

Tang, G., Peng, L., Baldwin, P. R., Mann, D. S., Jiang, W., Rees, I., and Ludtke, S. J. (2007). EMAN2: an extensible image processing suite for electron microscopy. J Struct Biol 157, 38–46.

Tang, L., Morris, J., Wan, J., Moore, C., Fujita, Y., Gillaspie, S., Aube, E., Nanda, J., Marques, M., Jangal, M., Anderson, A., Cox, C., Hiraishi, H., Dong, L., Saito, H., Singh, C. R., Witcher, M., Topisirovic, I., Qian, S. B., and Asano, K. (2017). Competition between translation initiation factor eIF5 and its mimic protein 5MP determines non-AUG initiation rate genome-wide. Nucleic Acids Res

Terenin, I. M., Akulich, K. A., Andreev, D. E., Polyanskaya, S. A., Shatsky, I. N., and Dmitriev, S. E. (2016). Sliding of a 43S ribosomal complex from the recognized AUG codon triggered by a delay in eIF2-bound GTP hydrolysis. Nucleic Acids Res 44, 1882–1893.

Thakur, A., and Hinnebusch, A. G. (2018). eIF1 Loop 2 interactions with Met-tRNA. Proc Natl Acad Sci U S A 115, E4159–E4168.

Thiaville, P. C., Legendre, R., Rojas-Benítez, D., Baudin-Baillieu, A., Hatin, I., Chalancon, G., Glavic, A., Namy, O., and de Crécy-Lagard, V. (2016). Global translational impacts of the loss of the tRNA modification t(6)A in yeast. Microb Cell 3, 29–45.

Valasek, L., Nielsen, K. H., Zhang, F., Fekete, C. A., and Hinnebusch, A. G. (2004). Interactions of eukaryotic translation initiation factor 3 (eIF3) subunit NIP1/c with eIF1 and eIF5 promote preinitiation complex assembly and regulate start codon selection. Mol Cell Biol 24, 9437–9455.

Valášek, L. S., Zeman, J., Wagner, S., Beznosková, P., Pavlíková, Z., Mohammad, M. P., Hronová, V., Herrmannová, A., Hashem, Y., and Gunišová, S. (2017). Embraced by eIF3: structural and functional insights into the roles of eIF3 across the translation cycle. Nucleic Acids Res

Walker, S. E., Zhou, F., Mitchell, S. F., Larson, V. S., Valasek, L., Hinnebusch, A. G., and Lorsch, J. R. (2013). Yeast eIF4B binds to the head of the 40S ribosomal subunit and promotes mRNA recruitment through its N-terminal and internal repeat domains. RNA 19, 191–207.

Wei, Z., Xue, Y., Xu, H., and Gong, W. (2006). Crystal structure of the C-terminal domain of S.cerevisiae eIF5. J Mol Biol 359, 1–9.

Yamamoto, H., Unbehaun, A., Loerke, J., Behrmann, E., Collier, M., Bürger, J., Mielke, T., and Spahn, C. M. (2014). Structure of the mammalian 80S initiation complex with initiation factor 5B on HCV-IRES RNA. Nat Struct Mol Biol 21, 721–727.

Zhang, K. (2016). Gctf: Real-time CTF determination and correction. J Struct Biol 193, 1–12.

Zur, H., and Tuller, T. (2013). New universal rules of eukaryotic translation initiation fidelity. PLoS Comput Biol 9, e1003136.

